# Unlocking TAS2R14 activation through intricate multi-ligand binding networks

**DOI:** 10.1101/2024.04.13.589249

**Authors:** Xiaolong Hu, Weizhen Ao, Mingxin Gao, Lijie Wu, Yuan Pei, Shenhui Liu, Qianqian Sun, Junlin Liu, Longquan Jiang, Yiran Wu, Xin Wang, Yan Li, Qiwen Tan, Jie Cheng, Fan Yang, Chi Yang, Jinpeng Sun, Tian Hua, Zhi-Jie Liu

**Affiliations:** iHuman Institute, ShanghaiTech University, Shanghai 201210, China; School of Life Science and Technology, ShanghaiTech University, Shanghai 201210, China; NHC Key Laboratory of Otorhinolaryngology, Qilu hospital and School of Basic Medical Sciences, Shandong University, Jinan, Shandong 250012, China; Department of Oral Surgery, Shanghai Ninth People’s Hospital and College of Stomatology, Shanghai Jiao Tong University School of Medicine; National Center for Stomatology and National Clinical Research Center for Oral Diseases, Shanghai Key Laboratory of Stomatology and Research Unit of Oral and Maxillofacial Regenerative Medicine, Chinese Academy of Medical Sciences

## Abstract

Bitter taste receptors, particularly TAS2R14, play central roles in discerning a wide array of bitter substances, ranging from dietary components to pharmaceutical agents^1–3^. In addition, TAS2R14 has broad expression in non-gustatory tissues, suggesting its important roles in selective physiological processes and therapeutic potential^4^. Here, we present cryo-electron microcopy structures of TAS2R14 in complex with flufenamic acid and aristolochic acid, coupling with G protein subtypes gustducin and G_i_. Distinct from most known GPCRs, agonists of TAS2R14 bind to multiple intracellular pockets. We highlight the cholesterol molecules occupying an upper transmembrane site, typically the orthosteric pocket in other GPCRs, also binding in the entrance to an intracellular agonist binding pocket, suggesting an endogenous modulatory function^5^. The structural and mutagenesis analysis illuminate the receptor’s broad-spectrum ligand recognition and activation via intricate multiple ligand-binding sites. Furthermore, we unveil the structural configuration of gustducin and its interaction with TAS2R14, as well as the coupling mode of G_i1_. This investigation should be instrumental in advancing our knowledge of the mechanisms underlying bitter taste recognition and response, with broader relevance to sensory biology and pharmacology.

## Introduction

G protein coupled-receptors (GPCRs) constitute the largest superfamily of membrane proteins, orchestrating a vast array of physiological functions^6^. Within this superfamily, taste receptors (TASRs) have specialized in discerning gustatory stimuli such as sweet (TAS1R), umami (TAS1R), and bitter (TAS2R) tastes^7,8^. In addition to their traditional gustatory roles, TAS2Rs are expressed in extraoral tissues, revealing a spectrum of potential functions that extend beyond bitter tasting^9,10^. Among the 25 TAS2Rs, TAS2R14 is a receptor of particular interest due to its promiscuity, responding to a variety of agonists with diverse structures^1,11^. Intriguingly, many clinical drugs evoke bitter taste responses, with approximately 9% of tested pharmaceutical drugs activating TAS2R14^2,3^. Moreover, TAS2R14 exhibits broad expression in extraoral tissues, including heart^12^ and airway smooth muscle (ASM)^4^. Notably, its activation in ASM induces bronchodilation, surpassing the effects triggered by β2-adrenergic receptor^13^. Consequently, TAS2R14 emerges as a potential therapeutic target for conditions like asthma^14^ or chronic obstructive pulmonary disease (COPD)^15^. For taste perception, TAS2Rs typically couple to the G protein gustducin^16,17^, leading to increased intracellular calcium. Of note, the structure of a native G protein gustducin coupled taste receptor is still illusive. In extraoral cells, the expression of Gα_gust_ is low, suggesting that TAS2Rs may instead couple with the prevalent G protein G_i_ for signal transduction^18^.

Among the known TAS2R14 agonists, flufenamic acid (FFA) and aristolochic acid (AA) are two well-characterized activators with suitable potency^1,2,19^. FFA, a nonsteroidal anti-inflammatory drug^20^, and AA, a potent nephrotoxin produced by *Aristolochia* plant^21^, are intriguing examples of diverse chemical structures capable of binding to TAS2R14. Moreover, the antitussive drug noscapine (NOS) and silibinin extracted from *milk thistle*^22^ were shown to activate TAS2R14^23^. In addition, cholesterol was reported to modulate the TAS2R14’s signaling in human airway cells^5^. The intriguing question arises: how do these therapeutic drugs and membrane cholesterol, each with distinct chemical structures, engage with one bitter taste receptor TAS2R14. Understanding the molecular mechanism of activation and signaling of TAS2R14 requires to unravel the complex interplay between these agonist drugs and the receptor. Thus, we present cryo-electron microscopy (cryo-EM) complex structures of TAS2R14 with AA or FFA, and wild type (WT) or chimeric gustducin, as well as WT G_i1_, elucidating the molecular details of ligand recognition, cholesterol modulation and different G protein coupling modes of this most promiscuous bitter taste receptor. This endeavor contributes valuable insights into the intricate pharmacology of TAS2R14, potentially informing novel therapeutic strategies.

## Results

### Structures of TAS2R14 in complex with gustducin and Gi

To enhance the stability of TAS2R14 complex samples for structural analysis, the WT human TAS2R14 was modified by introducing an N-terminal NanI sialidase (PDB code 2VK5) fusion protein, and assembled with chimeric gustducin, miniG_s/gust_, using the NanoBit strategy as described in TAS2R46 studies^24^ (Methods). Through extensive optimization processes, we further constituted TAS2R14 in complex with either WT Gustducin or G_i1_ (Fig. 1a, Extended Data Figs. 1-2). Based on previous reports, AA and FFA are two potent agonists for TAS2R14^25^. In our calcium mobilization assay, AA and FFA activated TAS2R14 with half maximum response (EC_50_) values of 10.3 and 10.4 µM, respectively (Fig. 1b). The varying concentrations for AA (50 and 150 µM) and FFA (150 µM to 1mM) were tested in the purification of TAS2R14-miniG_s/gust_ or TAS2R14-G_i_ complexes, separately. Eventually, we obtained the cryo-EM structures of 50 µM AA TAS2R14-G_i_ complexes, 150 µM AA TAS2R14-miniG_s/gust_ and TAS2R14-G_i_ complexes, at overall resolutions ranging from 2.7-3.1 Å (Fig. 1c, Extended Data Figs. 1, 3, Extended Data Table. 1), and the corresponding structures are referred as AA_50µM_-TAS2R14-G_i_, AA_150µM_-TAS2R14-miniG_s/gust_ and AA_150µM_-TAS2R14-G_i_, respectively. Using similar strategy, the cryo-EM structures of FFA_1mM_-TAS2R14-miniG_s/gust_ and FFA_350µM_-TAS2R14-G_i_ are determined at global resolutions of 3.2 and 3.3 Å, respectively (Fig. 1d, Extended Data Fig. 2a-b, Extended Data Table. 1).

**Fig. 1.**
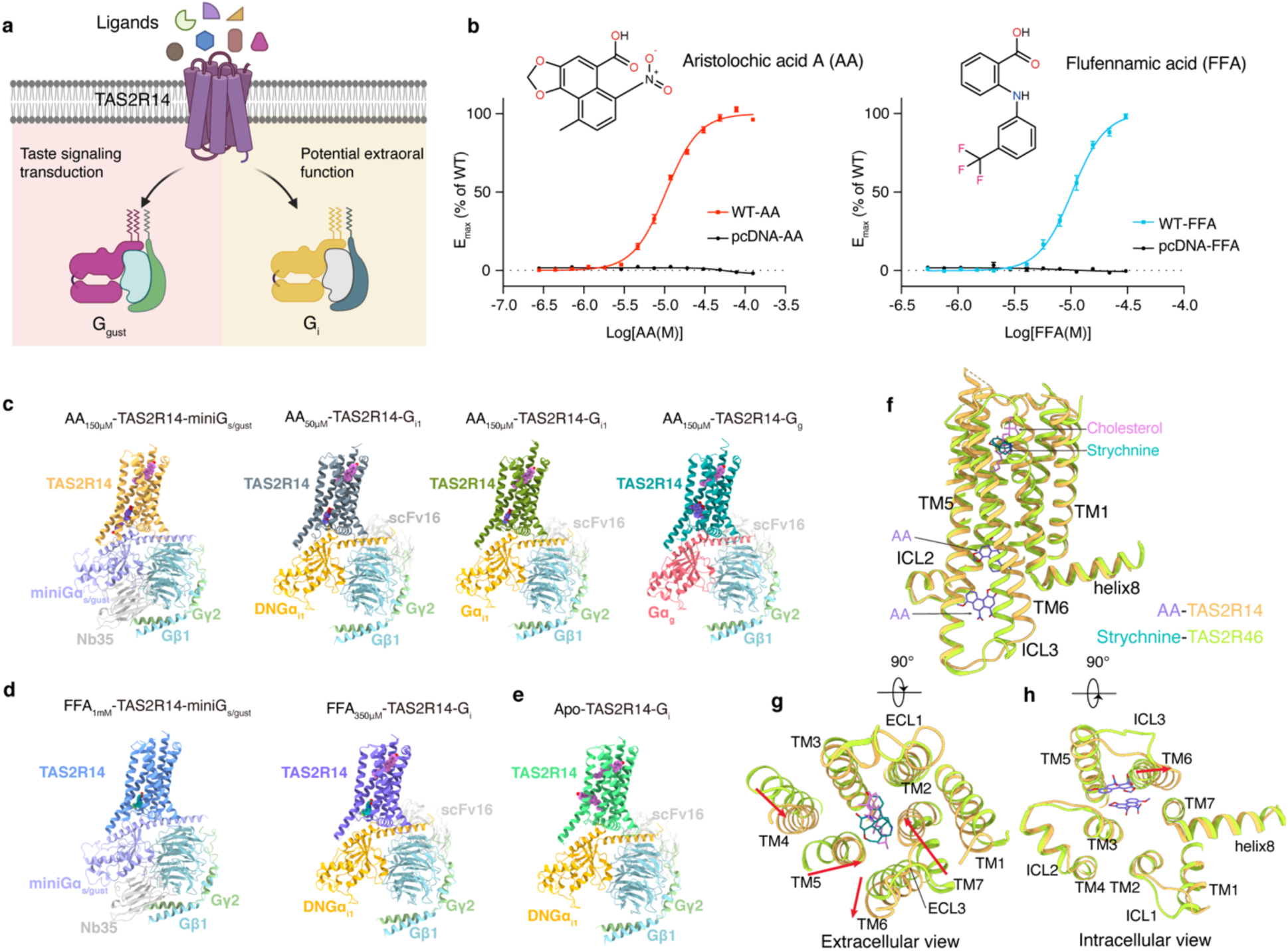
Cryo-EM structures of TAS2R14-G protein complexes. **a,** Schematic of TAS2R14-mediated activation of downstream gustducin and G_i_ signaling pathways by diverse ligands. **b,** Chemical structures of aristolochic acid A (AA) and flufennamic acid (FFA), and their activies on TAS2R14 measured in the FLIPR Ca^2+^assay. Data are mean ± s.e.m. of n ≥ 3 independent experiments performed in triplicate. **c,** Cryo-EM structures of AA_150µM_-TAS2R14-miniG_s/gust_, AA_50µM_-TAS2R14-G_i1_, AA_150µM_-TAS2R14-G_i1_ and AA_150µM_-TAS2R14-G_g_. **d,** Cryo-EM structures of FFA_1mM_-TAS2R14-miniG_s/gust_ and FFA_350µM_-TAS2R14-G_i1_. **e,** Cryo-EM structure of Apo-TAS2R14-G_i1_. **f-g,** Structural comparison of AA-TAS2R14 and strychnine-TAS2R46 in side view **(f)**, extracellular view **(g)** and intracellular view **(h)**.

Furthermore, the AA bound TAS2R14 in complex with WT gustducin was successfully solved after extensive optimization, referred as AA_150µM_-TAS2R14-G_g_, revealing the coupling modes of TAS2Rs with its native G protein (Fig. 1c, Extended Data Figs. 1d, 3e). Moreover, the cryo-EM structure of apo-TAS2R14-G_i_ was also determined at a global resolution of 2.9 Å, to facilitate comparative analysis with the agonist-bound conformations (Fig. 1e, Extended Data Fig. 2c, 3h). Comparative structural analysis reveals that TAS2R14 exhibits similar overall conformation across different ligand-bound states (Extended Data Fig. 4). The overall architecture of TAS2R14 closely resembles the previously elucidated TAS2R46^24^ (Fig. 1f-h), sharing 45% sequence identity and a RMSD of 1.4 Å, particularly in the intracellular domains, with the exception of transmembrane helix 6 (TM6) (Fig. 1f). However, notable differences were observed in the extracellular regions, including the extracellular loops (ECLs) and ligand-binding pockets (Fig. 1g). Specifically, TAS2R14’s extracellular domain is comparatively more compact, and its TM6 exhibits a higher degree of flexibility relative to TAS2R46 (Fig. 1g-h).

### Cholesterol and multiple AA binding pockets in TAS2R14

TAS2R14, a member of the broadly turned bitter taste receptor category, exhibits a minimal preference for structural features of subsances^25^. Through cryo-EM structural analysis, we identified three ligand binding sites indicated by EM density maps in TAS2R14 complexed with miniG_s/gust_ and G_i1_ (Fig. 2a, Extended Data Fig. 3b-g). Specifically, the AA_150µM_-TAS2R14-miniG_s/gust_ structure reveals the presence of a cholesterol molecule in pocket-1 and two AA molecules in pockets 2 and 3 (Fig. 2a, Extended Data Fig. 3b), suggesting a complex ligand interaction scheme.

**Fig. 2.**
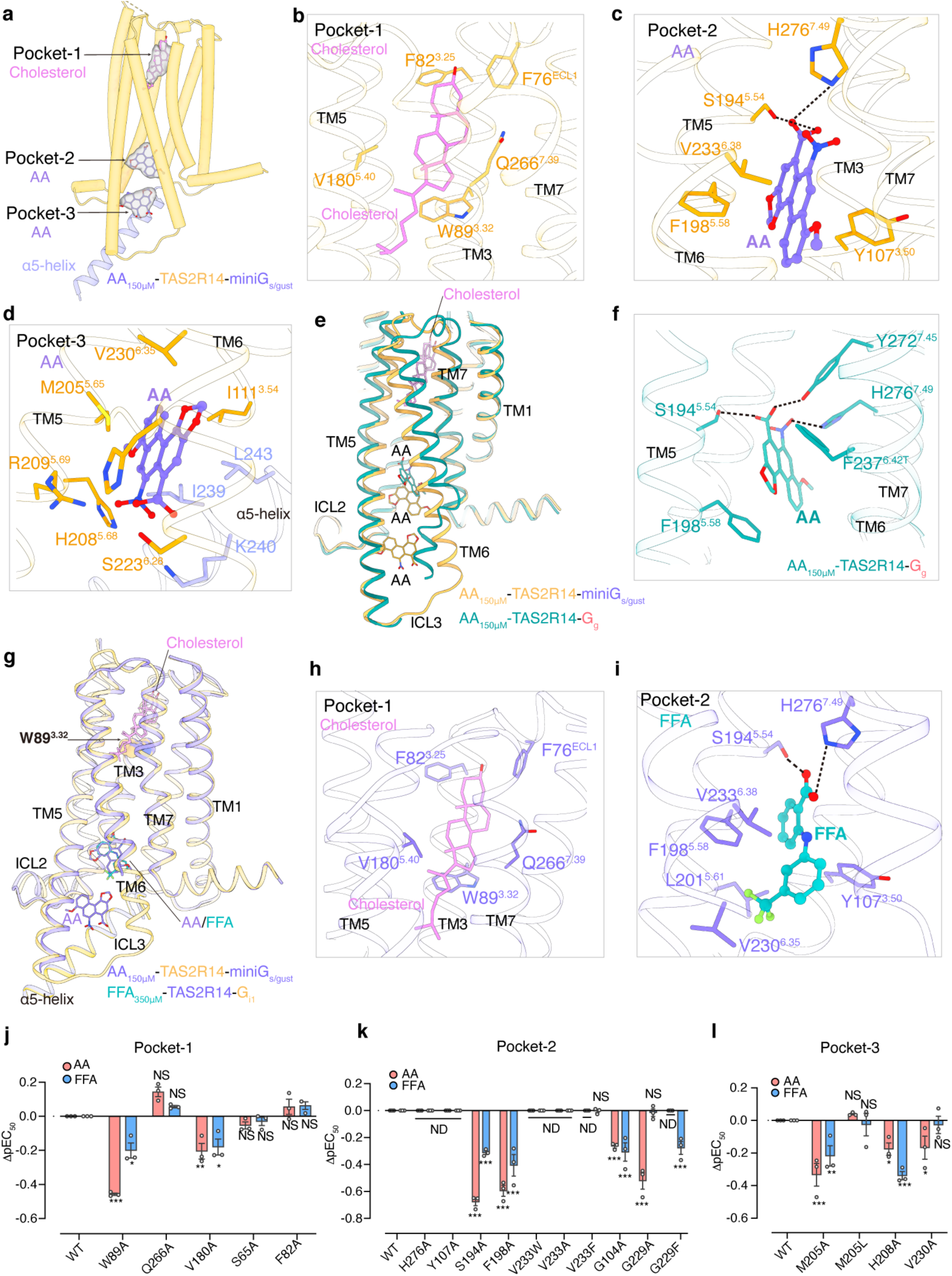
Recognition mechanism of AA, FFA and cholesterol by TAS2R14. **a,** Three binding pockets in AA_150µM_-TAS2R14-miniG_s/gust_ complex structure. The receptor is shown as tube cartoon, the cryo-EM densities for cholesterol and each AA are shown as gray surfaces. **b,** Cholesterol binding in pocket-1. **c-d,** Key residues in the AA binding pocket-2 **(c)** and pocket-3 **(d)** in TAS2R14. e, Structure superposition of binding poses of cholesterol and AA in AA_150µM_-TAS2R14-miniG_s/gust_ and AA_150µM_-TAS2R14-G_g_ complexes. **f,** AA binding pose in AA_150µM_-TAS2R14-G_g_ structure and key residues are shown as sticks. **g,** Structure superposition of binding poses of AA and FFA in AA_150µM_-TAS2R14-miniG_s/gust_ and FFA_350µM_-TAS2R14-G_i1_ complexes. **h,** Cholesterol binding pose in FFA_350µM_-TAS2R14-G_i1_ structure. **i,** Key residues for FFA binding in pocket-2 in TAS2R14. **j-l,** FLIPR Ca^2+^assay responses in wildtype (WT) TAS2R14 for mutants in pockets 1, 2 and 3 following AA or FFA stimulation. The negative logarithmic half-maximal effective concentration (pEC_50_) values were calculated from the concentration–response curves (Extended Data Fig. 5). *P < 0.05; **P < 0.01; ***P < 0.001, calculated using one-way analysis of variance (ANOVA) followed by Dunnett’s test for multiple comparison analysis (with reference to the WT). NS, not significantly different between the groups. ND, signal not detectable

Pocket-1 overlaps with the conventional orthosteric pocket, composed of extracellular portions of TMs 2, 3, 5 and 7 (Fig. 2b), where cholesterol molecule engages the conserved residue W89^3.32^ through π-π interactions and establishes hydrophobic interactions within the pocket in AA_150µM_-TAS2R14-miniG_s/gust_ structure (Fig. 2b). The binding of cholesterol, not AA, is also observed in the pocket-1 of apo-TAS2R14-G_i_ structure (Extended Data Fig. 3h), indicating cholesterol’s potential role as an endogenous modulator of TAS2R14. Consistently, only the alanine mutation of the conserved residue W89^3.32^ significantly reduces AA’s activity toward the receptor (Fig. 2j), single mutations of other residues in pocket-1 have a lesser impact in our calcium mobilization assay (Fig. 2j, Extended Data Fig. 5a,d,g, Extended Data Table 2). The mutagenesis evidence supports the hypothesized role of cholesterol and refines our understanding of ligand interactions within pocket-1 of TAS2R14.

Pocket-2 resides within the intracellular half of 7TM bundle and is formed by TMs 3, 5-7 (Fig. 2c). AA can be unambiguously modeled into the EM density in this pocket (Extended Data Fig. 3b-e). Here, AA primarily establishes hydrophobic contacts or π-π interactions with residues Y107^3.50^, F198^5.58^ and V233^6.38^. Additionally, AA’s carboxylic acid group establishes hydrogen bonds with H276^7.49^ and S194^5.54^ (Fig. 2c). Mutations of S194^5.54^A, F198^5.58^A or V233^6.38^A significantly reduce the AA’s activity on TAS2R14 (Fig. 2k, Extended Data Fig. 5b,e,h). Y107^3.50^ and H276^7.49^ are two conserved residues in TAS2Rs, mutation of Y107^3.50^A or H276^7.49^A abolishes the receptor activation induced by AA (Fig. 2k, Extended Data Fig. 5l).

Pocket-3 is situated bellow pocket-2, shaped by the cytoplasmic ends of TM5 and TM6, and the α5 helix of miniGα_s/gust_ (Fig. 2d). AA’s interaction with pocket-3 involves hydrophobic interactions with residues I111^3.54^, M205^5.65^, V230^6.35^ from receptor and L243 and I239 from the α5 helix, along with polar interactions with H208^5.68^, R209^5.69^ and H227^6.32^ (Fig. 2d). Notably, M205^5.65^ is pivotal for AA binding to pocket-3, as evidenced by a conformation change upon AA binding (Extended Data Fig. 4b). This residue is unique to TAS2R14, while other TAS2Rs have Leu at this position. M205^5.65^A mutation affects both the efficacy and potency of AA on TAS2R14, while M205^5.65^L mutation does not (Fig. 2l, Extended Data Fig. 5l), indicating the bulky hydrophobic residue is important for either ligand binding or stabilizing the conformation in this region. The presence of multiple binding sites of AA and cholesterol in TAS2R14 suggests an unusual mechanism for its recognition of a broad-spectrum of ligands with diverse scaffolds, potentially through a multifaceted activation mechanism.

For the G_i_-coupled structures, AA_50µM_-TAS2R14-G_i_ and AA_150µM_-TAS2R14-G_i_, cholesterol binds in pocket-1 and AA occupies pocket-2, similar to those in TAS2R14-miniG_s/gust_ complexes (Extended Data Fig. 4a-b). However, the cytoplasmic end of TM6 (D221^6.26^-R228^6.33^), becomes more ordered and bends towards TM5 in G_i_-coupled TAS2R14 structures (Extended Data Fig. 4a), interfering with ligand binding at pocket-3. In the cryo-EM structure of AA_150µM_ -TAS2R14-G_g_ complex, while the receptor and gustducin are clearly defined, the EM density for cholesterol in pocket-1 is weaker than that in miniG_s/gust_- or G_i_-coupled complex structures (Extended Data Fig. 3e). However, an AA molecule binds at the entrance to pocket-2 and partially overlaps with cholesterol’s binding spot in apo-TAS2R14-G_i_ structure (Fig. 2e-f). It forms hydrogen bonds with H276^7.49^ and S194^5.54^ and π-π interactions with residues F198^5.58^ and F237^6.42T^ (Fig. 2f). The bound AA displays altered binding pose comparing with AA in AA_150µM_-TAS2R14-miniG_s/gust_ or AA_150µM_-TAS2R14-G_i_ structures, potentially activates the receptor. The structure differences, including the altered AA binding, suggest that G protein coupling may allosterically modulate the interaction between receptor and ligands.

### FFA binding in TAS2R14

Similar to the observations in AA bound TAS2R14 structures (Fig. 2a), in FFA_350µM_-TAS2R14-G_i_ complex structure, the EM densities for cholesterol and anti-inflammatory drug FFA were also identified in pockets 1 and 2, respectively, except that the EM density for cholesterol is comparatively weaker (Fig. 2g, Extended Data Fig. 3g). The cholesterol is located within pocket-1, forming π-π interaction with W89^3.32^ (Fig. 2h), and the mutation of W89^3.32^A also affects FFA’s activity toward TAS2R14, indicating the role of W89^3.32^ in stabilizing the conformation of this region. Accordingly, single mutations of other residues in pocket-1 also slightly affect FFA-induced activation of TAS2R14 in both calcium mobilization and G protein dissociation assays (Fig. 2j, Extended Data Figs. 5a,d, 6a). In pocket-2, FFA establishes strong interactions with the receptor, including hydrogen bonds between the carboxylic acid group of FFA and H276^7.49^ and S194^5.54^ that mirror those observed with AA (Fig. 2i). FFA’s trifluoromethyl group additionally forms hydrophobic interactions with I111^3.54^, L201^5.61^, I202^5.62^ and V230^6.35^ (Fig. 2i). Furthermore, the diphenylamine core of FFA forms hydrophobic and π-π interactions with Y107^3.50^ and F198^5.58^ (Fig. 2i). Consistent with these observations, alanine mutations of S194^5.54^, Y107^3.50^ and H276^7.49^ significantly affect FFA’s activation on TAS2R14 in both calcium mobilization and G protein dissociation assays (Fig. 2k, Extended Data Figs. 5b,e, Extended Data Fig. 6b).

In FFA_1mM_-TAS2R14-miniG_s/gust_ structure, FFA occupies pocket-2 and adopts a binding pose similar to that in the G_i_-coupled FFA-TAS2R14 structure (Extended Data Fig. 4c). However, the EM density in pocket-1 is notably weaker (Extended Data Fig. 3f), which prevents the accurate modeling of the cholesterol molecule. These observations indicate probable cross-modulations between agonist and cholesterol in TAS2R14 signaling.

A noted observation from the structural and ligand-binding analysis is the highly dynamic nature of TM6 and its role in coordinating ligand binding to the intracellular pockets of TAS2R14 (Extended Data Fig. 4 and 7a). In the apo-TAS2R14-G_i_ complex structure, TM6 (S232^6.37^-D221^6.26^) exhibits flexibility, indicated by the absence of EM density (Extended Data Fig. 2c). Conversely, when the ligand binds to intracellular pockets 2 and 3, the ordered TM6 structure extends into the cytosol region, indicated by the clear EM density (Extended Data Figs. 3c-h and 7a). The adaptive behavior of TM6 is indicative of its special function in modulating the receptor’s binding capacity, providing new insights into the understanding of the unique features in bitter taste receptors.

### Characterization of ligands binding pockets in TAS2R14

Structure analysis of TAS2R14 has uncovered three discrete binding sites–another novel character that improve our understanding of ligand recognition, cholesterol modulation and receptor activation mechanisms in TAS2Rs. Of these, cholesterol binds in pocket-1 – the only pocket opening to the extracellular side and overlapping with the canonical orthosteric binding pocket in most class A GPCRs. This pocket is anchored by the conserved Trp^3.32^ as observed in TAS2R46 structure^24^ (Extended Data Fig.7b). Furthermore, no other individual mutations of residues within pocket-1 results in a complete loss of receptor function for either AA or FFA (Fig. 2j). Of note, a second cholesterol molecule was also identified in the cleft between TM5 and TM6 in the intracellular region formed by residues F191^5.51^, F198^5.58^ and F237^6.42T^, positioned at the entry path to pocket-2 (Extended Data Figs.3h and 7a). Those findings hint cholesterol’s allosteric modulation role in ligand binding and activation in TAS2R14 through both pocket-1 and pocket-2.

The pocket-2 is nestled in the receptor’s intracellular core. Our sequence analysis of 25 TAS2Rs, highlights that residues G104^3.47^ and G229^6.34^, involved in the constitution of pocket-2, are exclusive to TAS2R14 (Extended Data Fig.5l). These unique residues likely facilitate the pocket with necessary plasticity and capacity to accommodate diverse ligands. Concordantly, alanine mutations of G104^3.47^ or G229^6.34^ markedly reduce AA- or FFA-induced activity, and the G229^6.34^F mutation almost abolishes TAS2R14’s activity (Fig. 2k, Extended Data Figs. 5 and 6b). Moreover, pocket-2 is in proximity to activation-related motifs, HS/P^7.50^FIL and FY^3.50^L, as previously identified for TAS2Rs^24,26^. Thus, agonist binding in pocket-2 is crucial for activation and active conformation stabilization of TAS2R14 (Fig. 3a). As mentioned above, either Y107^3.50^A or H276^7.49^A mutation abolishes or significantly affects the activity of AA or FFA on TAS2R14 (Fig. 2k, Extended Data Figs. 5 and 6b). Additionally, V233^6.38^ forms hydrophobic interactions with phenanthrene ring of AA or diphenylamine ring of FFA (Fig. 2g, i). V233^6.38^A mutation impairs both AA- and FFA-induced activity on TAS2R14, however, V233^6.38^F selectively disrupts AA-induced activity (Fig. 2k), probably due to the steric clashes with AA’s bulkier structure. Collectively, pocket-2 is vital for ligand-induced activation of TAS2R14.

**Fig. 3.**
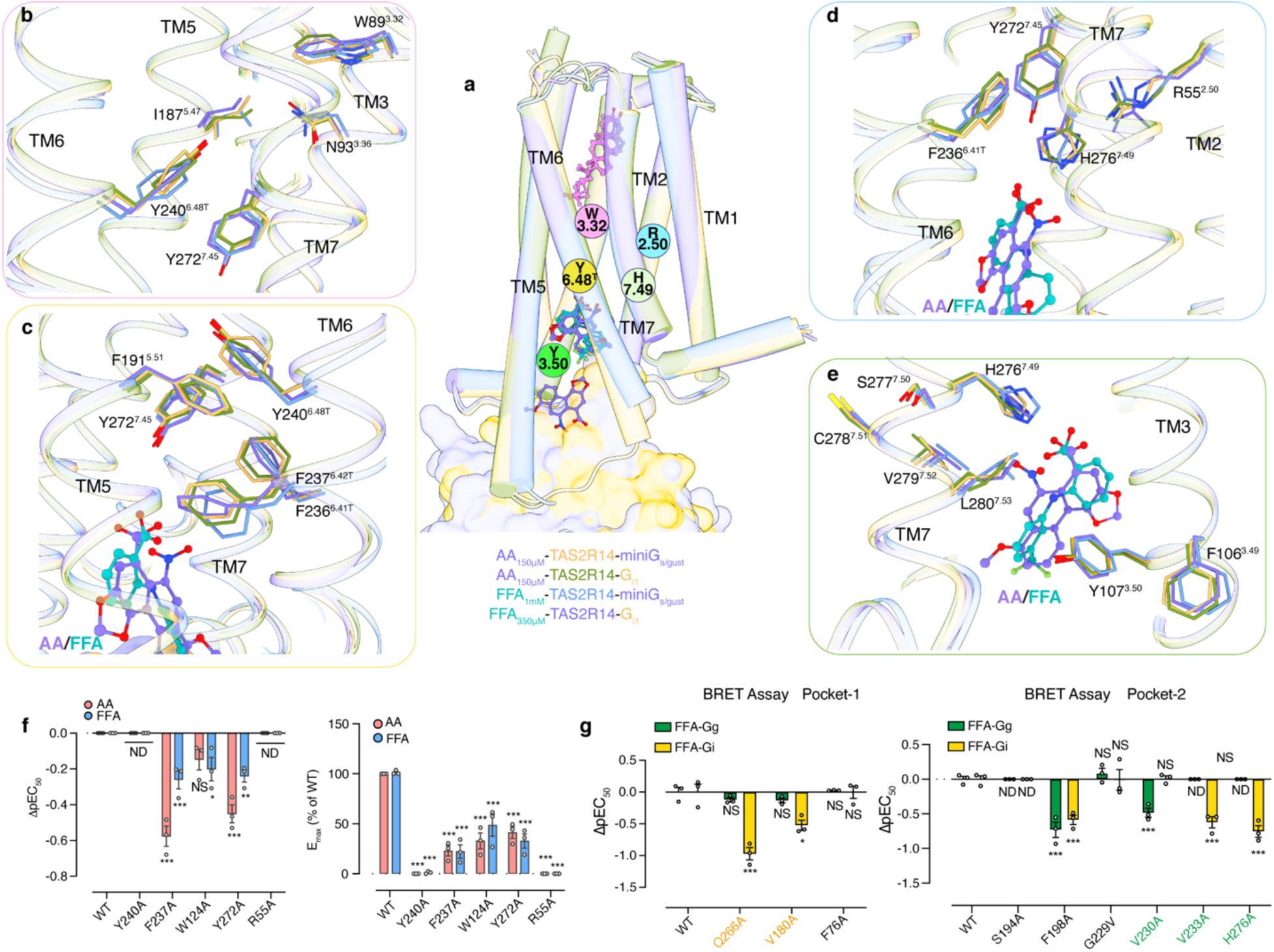
The activation features of TAS2R14. **a,** Schematic presentation of the key residues that contribute to TAS2R14 in active state. AA_150µM_-TAS2R14-miniG_s/gust_, AA_150µM_-TAS2R14-G_i1_, FFA_1mM_-TAS2R14-miniG_s/gust_ and FFA_350µM_-TAS2R14-G_i1_ complex structures are used for analysis. **b-e,** The detailed interactions of residues that stabilize the conformations of W89^3.32^ and Y240^6.48T^ region **(b)**, TM5-TM6-TM7 core **(c)**, network formed by R55^2.50^, Y272^7.45^ and H276^7.49^ **(d)** and networks formed by HS/P^7.50^FIL, FY^3.50^L motifs and ligands in pocket-2 **(e)**. **f,** FLIPR Ca^2+^assay responses in wildtype (WT) TAS2R14 for mutants related to the receptor activation following AA or FFA stimulation. The negative logarithmic half-maximal effective concentration (pEC_50_) and the maximum response (*E*_max_) values were calculated from the concentration–response curves (Extended Data Fig. 5). **g,** Effects of key residue mutations in pockets 1 and 2 on TAS2R14’s coupling with G_g_ or G_i1_ induced by FFA in BRET assay. Data are mean ± s.e.m. from 3 independent experiments (n = 3). Two-sided one-way ANOVA with Tukey’s test (compared with wild type). *P < 0.05; **P < 0.01; ***P < 0.001; NS, not significant. ND, signal not detectable.

The pocket-3 is occupied only in the AA_150µM_-TAS2R14-miniG_s/gust_ structure, the outward rotation of M205^5.65^ facilitates AA’s binding in this site (Extended Data Fig. 4b). Here, AA functions as a molecular wedge, stabilizing the intracellular parts of TM5, TM6 and the α5 helix of G protein (Fig. 2d). The ligand could act either as a positive allosteric modulator (PAM), stabilizing the active conformation of TM6, or as a negative allosteric modulator (NAM), slowing down the dissociation of Gα from the receptor. This dual potential underscore the need for further research to clarify AA’s modulatory role and its implications for the signaling dynamics of TAS2R14.

### Activation features of TAS2R14

To elucidate the activation mechanism of TAS2R14, we performed a comparative structural analysis using active and apo TAS2R14 forms (Extended Data Fig. 7a). Notably, the most significant conformational changes among those structures occur in TM6 (Extended Data Fig. 7a), along with subtle changes in TM5 and TM7. Specifically, TM6 is ordered only in the extracellular portion, when cholesterol bind to pocket-1 or in the absence of agonists. As agonists occupy pockets 2 and 3, the ordered structure of TM6 extends into the cytosol domain. In this process, TM6 acts as a “zipper” moving down and sealing the exposed ligand binding pocket-2 and 3 along the intracellular 7TM bundle (Extended Data Fig. 7a). In addition, residue Y240^6.48T^, corresponding to “toggle switch” residue Y241^6.48^ in TAS2R46 structures^24^, packs closer to the transmembrane bundle core upon ligand binding into pocket-2 (Fig. 3b). The Y240^6.48T^A mutation abolishes AA or FFA induced activation of TAS2R14 in our calcium mobilization assay (Fig. 3f). With the binding of AA or FFA into pocket-2, F237^6.42T^ rotates outward, forming hydrophobic and π-π interactions with residues from TM5 (Fig. 3c). The F237^6.42T^A mutation affects the activities of AA and FFA on TAS2R14 (Fig. 3f). Ligand engagement in pocket-2 also helps to stabilize TM3, 5, 6 and 7 bundle through interactions with the conserved motifs HS/P^7.50^FIL (H276^7.49^S277^7.50^C278^7.51^V279^7.52^L280^7.53^ in TAS2R14) and FY^3.50^L (F106^3.49^Y107^3.50^L108^3.51^ in TAS2R14) (Fig. 3d-e). These findings reveal a distinct role of TM6 in coordinating the ligand binding and receptor activation of TAS2R14, in addition to its conventional function in G protein coupling to GPCRs. We next exploited G_g_ and G_i1_ protein dissociation assays in parallel to investigate the role of key residues within TAS2R14’s ligand-binding pockets and their response to FFA activation through G_g_ and G_i1_ pathways (Fig. 3g). The results are consistent with that in the calcium mobilization assay. Mutations of pocket 2 may have effects on either G_i_ or G_g_ coupling (Fig. 3g). The V230^6.35^A mutation has more effects on G_g_ coupling than G_i1_ coupling. These results suggested the possibility of distinct activation mechanisms in TAS2R14 for individual G protein pathway.

Comparing the active TAS2R14 and TAS2R46 structures, the conserved activation-related motifs, R^2.50^, Y^6.48^, HS/P^7.50^FIL and FY^3.50^L in class T GPCRs, adopt similar conformations, as well as the strictly conserved residue Trp^3.41^ which anchors the spatial arrangement of TM4 and TM5 (Extended Data Fig. 7b-d). Additionally, the opening angle of the cytoplasmic parts of TM5 and TM6 in TAS2R14 is larger than in TAS2R46 (Fig. 1f), due to the extra ligand binding pocket within this region in TAS2R14. These structural correlations contribute to a more profound comprehension of the shared and unique activation mechanisms among TAS2Rs, thereby enhancing our understanding of the diverse functionalities of these receptors.

### Gustducin and G_i_ coupling in TAS2R14

Gustducin (G_g_) is the endogenous transducer of taste receptors. The TAS2R14-G_g_ complex structure reveals the binding mode between a bitter taste receptor and gustducin (Fig. 4a). Of note, the TAS2R14-Gα_g_ binding mode is similar to that of TAS2R14-miniGα_s/gust_ (Extended Data Fig. 7e), suggesting the chimeric G protein is valid to mimic the coupling between native gustducin and the receptor. The interactions between TAS2R14 and G_g_ primarily occur through the intracellular regions of the receptor (TM2, TM3, ICL2, TM5, TM7 and helix 8) and the α5 and αN helices of the Gα_g_ subunit (Fig. 4c-d). Specifically, Y107^3.50^ forms hydrophobic interactions with L353 and C351 in Gα_g_ (Fig. 4c). Additionally, Gα_g_-L348 and I344, in conjunction with I111^3.54^ and M205^5.65^ from the receptor, contribute to the hydrophobic core, stabilizing the interaction interface (Fig. 4c). Notably, the conserved residue H208^5.68^ in TAS2Rs forms a hydrogen bond with Gα_g_-D341 in TAS2R14-G_g_ complex structure (Fig. 4c). Similar to TAS2R46, the ICL2 of TAS2R14 also forms a short helix, orienting parallel to the cell membrane (Fig. 4d). N113^ICL2^ points to the polar interaction core formed by residues K207^5.67^ and H208^5.68^ from TM5 of TAS2R14 (Fig. 4d). Furthermore, W124^ICL2^ participates in the interactions with the αN helix of Gα_g_ (Fig. 4d). Notably, the conserved residue at this site among TAS2Rs is positively charged residue. The H208^5.68^A or W124^ICL2^A mutation significantly affects the activity of TAS2R14 (Fig. 2l, 3f).

**Fig. 4.**
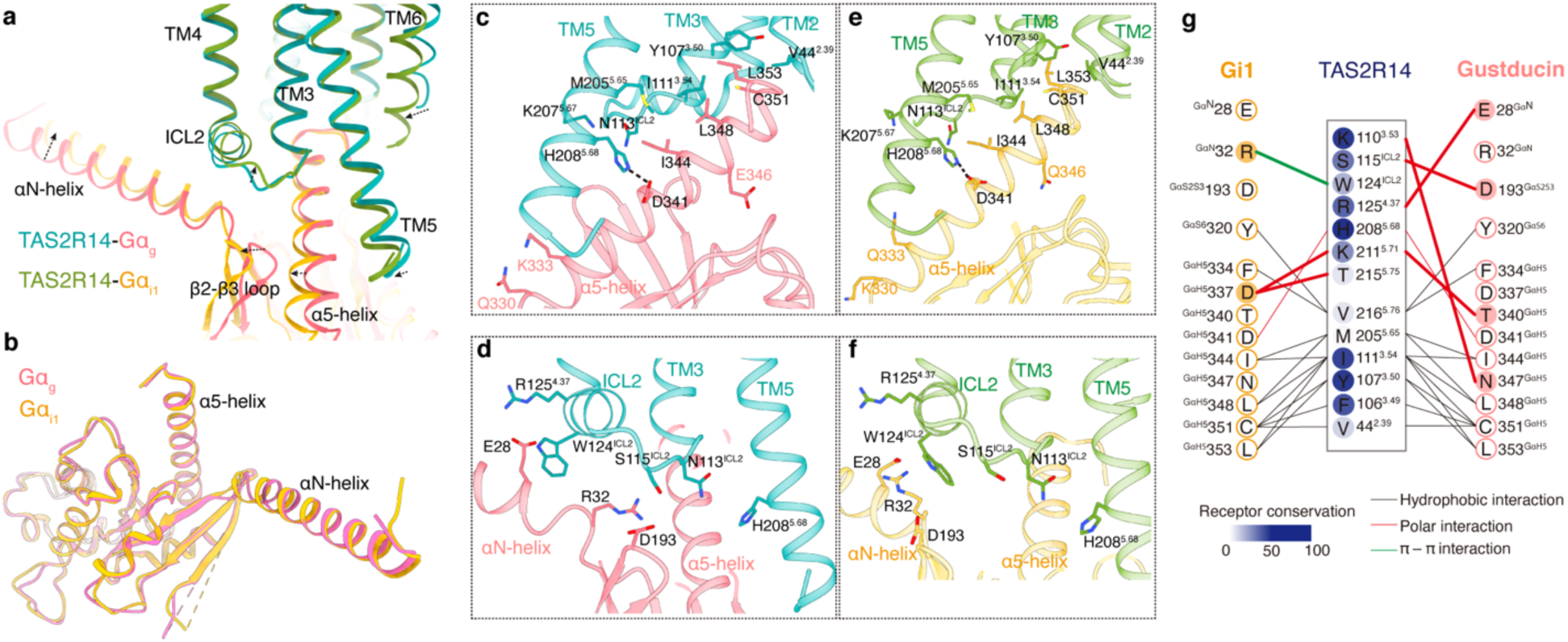
Gustducin and Gi coupling modes in TAS2R14. **a,** Structure comparison of the gustducin and G_i1_ binding modes in TAS2R14. AA_150µM_-TAS2R14-G_g_ and AA_150µM_-TAS2R14-G_i1_ complex structures are used for analysis. **b,** Structure alignment of Gα_g_ and Gα_i1_ subunits. **c-d,** The detailed interactions between TAS2R14 and Gα_g_ binding interface. **e-f,** The detailed interactions between TAS2R14 and Gα_i1_ binding interface. **g,** Diagram of the contacts between Gα_g_ and TAS2R14, and Gα_i1_ and TAS2R14, respectively.

In comparison with other TAS2Rs, TAS2R14 expresses in several extraoral tissues, where gustducin may not be the dominant G protein, G_i_ may instead contribute to the signal transductions^18^. In our BRET assays, FFA activated TAS2R14 shows coupling with Gα_i1_ (Fig. 3g). The structures of AA-TAS2R14-G_i1_ and FFA-TAS2R14-G_i1_ provide examples of a bitter taste receptor coupling with G_i_ protein (Fig. 1c-d), which show notable differences with G_i_ coupling mode in class A GPCRs, such as CB1 (Extended Data Fig. 7f). When comparing the coupling mode between TAS2R14 and Gα_g_, the Gα_i1_ subunit adopts a similar binding mode and interactions with TAS2R14 (Fig. 4a-b). Based on the sequence alignment of Gα_i1_ and Gα_g_, only four different residues are present in the α5 helix (Extended Data Fig. 3a), where E/N346 and Q/K330 are not involved in the interactions with the receptor, only Gα_g_-K333 and Gα_i1_-Q333 make different interactions with the cytoplasmic end of TM5 (Fig. 4c, 4e). However, the C-terminal end of αN helix in Gα_g_ or Gα_i1_ forms interactions with the receptor’s ICL2 in TAS2R14, while the sequences in the αN helix are more diverse in Gα_i1_ and Gα_g_ compared with that in α5 helix (Fig. 4d. 4f, 4g). These observations suggest that ICL2 may play a role in mediating TAS2Rs to couple with different G protein subtypes.

## Discussion

The present study offers a comprehensive exploration of TAS2R14, a taste receptor, and its interaction with different bitter compounds, including clinical drugs. The structural analysis reveals the receptor’s sophisticated mechanism for ligand binding, characterized by its adaptability across three distinct binding sites. The findings highlight the unique role of cholesterol in allosteric modulation from the canonical orthosteric pocket, the significant activation function of agonists in the unconventional intracellular pocket-2, and the possibility of PAM or NAM modulation by agonists in pocket-3, located near the cytosol region. This multifaceted approach enables TAS2R14 to recognize and respond to a wide range of ligands.

This study also elucidates the coupling modes of TAS2R14 with different G protein subtypes, specifically WT gustducin and G_i_. Moreover, the comparison between the complex structures of TAS2R14 at different states uncovers the pivotal role of TM6 in modulating ligand engagement to the intracellular regions of the receptor and G protein subtype coupling. These insights offer a deeper understanding of taste perception mechanisms and pave the way for novel pharmacological and therapeutic strategies targeting TAS2R14 and related receptors.

## Methods

### Cell lines

*Spodoptera frugiperda* (Sf9, Expression systems) and *Trichoplusia ni.* (High Five, Thermo Fisher) cells were gown in ESF medium at 27°C and 120 rpm. The HEK293T (ATCC, CRL-11268) cells were grown in a humidified 37°C incubator with 5% CO_2_ and maintained in Dulbecco’s Modified Eagle Medium (DMEM, Gibco) containing 10% fetal bovine serum (FBS, Gibco).

### Constructs

Human TAS2R14 (residues G2-S317) gene sequence was synthesized and cloned into pFastBac vector. To facilitate expression and purification, a haemagglutinin-signaling peptide (HA) sequence followed by a Flag tag, a 10×His tag, a TEV protease cleavage site and a fusion protein comprising the catalytic domain of *C. perfringens* NanI sialidase (PDB: 2VK5). To increase the uniformity and stability of the complexes, the NanoBiT strategy was employed^27,28^, in which the C terminus of TAS2R14 was fused to the large part of NanoBit (LgBiT), and the Gβ_1_ subunit’s C-terminus was fused to the small part of NanoBit (SmBit). For the miniGα_s/gust_, miniGα_s_ was used as the body, and the α5 helix was replaced with the sequence of the Gα_gust_ subunit (residues 329-354) to form a miniGα_s/gust_, analogous to that in TAS2R46^24^. In the TAS2R14-G_i_ complex, the previously reported G203A and A326S mutations were incorporated into the wild type (WT) Gα_i1_ subunit, resulting in DNGα_i1_ (dominant negative, DN)^29^. In the TAS2R14-G_g_ complex, 14 single point mutations were made in the αN helix of Gα_gustducin_ to form Gα_g-mutations_ (abbreviated as Gg). For functional studies, WT TAS2R14 (residues M1-S317) was subcloned into the pcDNA3.1 vector, with a Flag tag added to the receptor’s N-terminus. To enhance the expression of the receptor, 45 amino acids from the N-terminus of rat somatostatin receptor type 3^30^ were inserted between the Flag tag and TAS2R14. Moreover, the C-terminal 44 amino acids of Gα_16_ were replaced with the corresponding 44 amino acids from the C-terminal end of Gα_gust_ (residues 311-354) to create a Gα_16/gust44_ chimera for calcium mobilization functional assays^31^.

### Preparation of Nb35 and scFv16 antibodies

Nanobody-35 (Nb35) was expressed in *Escherichia coli* BL21 and purified as previously described^32^. In brief, Nb35 was purified using Ni-NTA affinity resin and then loaded into a size exclusion Superdex 75 Increase 10/300 GL column (Cytiva) equilibrated in a buffer containing 20 mM HEPES (pH7.5) and 100 mM NaCl, and stored at -80 °C for subsequent use.

Single-chain antibody scFv16 was expressed in High Five insect cells (Invitrogen) using the Bac-to-Bac baculovirus expression system. A GP67 signal peptide was added to the N-terminus of scFv16, while a TEV cleavage site and an 8×His tag were fused to its C-terminus. Infected cells were harvested by centrifugation after 48 h of incubation at 27°C. scFv16 was purified as previously described^33^. scFv16 was also purified using Ni-NTA affinity resin and then loaded into a size exclusion column (Superdex 75 Increase 10/300 GL, Cytiva) equilibrated in a buffer containing 20 mM HEPES (pH7.5) and 100 mM NaCl. The purified protein was stored at -80°C for further use.

### Expression and purification of TAS2R14-G protein complex

TAS2R14, miniGα_s/gust_ or DNGα_i1_ or Gα_g_, Gβ_1_γ_2_ were co-expressed in *Sf*9 insect cells (Invitrogen) at a 6:1:2 ratio for each virus using the Bac-to-Bac baculovirus expression system (Thermo Fisher). Cell cultures were grown in ESF 921 medium (Expression Systems) and harvested after 48 h of incubation at 27°C before collection by centrifugation, and the cell pellets were stored at -80°C for future use. Cell pellets were lysed with buffer 50 mM HEPES pH7.5, 50 mM NaCl, 2 mM MgCl_2_, and EDTA-free protease inhibitor cocktail (Roche). The TAS2R14-G protein complex was formed in membranes by the addition of 4 µg/ml Nb35 or 6 µg/ml scFv16, 1.25 U apyrase (NEB), 300 mM TCEP, 50 μM, 150 μM, 250 μM Aristolochic acid A (AA, MedChemExpress) or 150 μM, 350 μM, 1 mM Flufenamic acid (FFA, MedChemExpress), or without ligands, incubation for 4 h at room temperature. The raw membranes were collected by ultracentrifugation and then were solubilized with buffer 50 mM HEPES pH 7.5, 100 mM NaCl, 1% (w/v) lauryl maltose neopentylglycol (LMNG, Anatrace), 0.2% (w/v) cholesteryl hemisuccinate TRIS salt (CHS, Sigma-Aldrich), 1.25 U apyrase (NEB), 300 mM TCEP, 4 µg/ml Nb35 or 6 µg/ml scFv16 for 3 h at 4°C, immediately followed by ultracentrifugation at 35,000 rpm for 30 min at 4°C, and the supernatant was incubated with TALON IMAC resin (Clontech) and 20 mM imidazole for 12 h at 4°C. The resin was washed with 15 CVs of Wash Buffer I containing 25 mM HEPES pH7.5, 100 mM NaCl, 0.05% LMNG, 0.01% CHS, 30 mM imidazole, 10% glycerol and corresponding concentrations of AA or FFA, and 15 CVs of Wash Buffer II 25 mM HEPES pH 7. 5, 100 mM NaCl, 0.01% LMNG, 0.002% CHS, 30 mM imidazole, 10% glycerol and corresponding concentrations of AA or FFA. Complex was eluted with 4 CVs of elution buffer 25 mM HEPES pH 7.5, 100 mM NaCl, 0.01% LMNG, 0. 002% CHS, 250 mM imidazole, 10% glycerol and corresponding concentrations of AA or FFA. The sample was further purified by size-exclusion chromatography on a Superose 6 or Superdex 200 increase 10/ 300 GL column equilibrated in buffer containing 25 mM HEPES pH7.5, 100 mM NaCl, 300 μM TCEP, 0.00075% (w/v) LMNG, 0.00015% (w/v) CHS, 0.00025% (w/v) glyco-diosgenin (GDN, Anatrace) and corresponding concentrations of AA or FFA. The peak fractions, which contain the complex, were pooled and concentrated to 4-12 mg/ml for cryo-EM sample preparation. For structure determination of AA-TAS2R14-G_g_ and apo-TAS2R14-G_i_ complex, the purification followed the same protocol as mentioned above.

### Cryo-EM grid preparation and data collection

For cryo-EM grid preparation, samples for the different complexes at a concentration of 4-12 mg/ml. 3 μl sample were applied individually to electron microscopy grids (Quantifoil, 400 mesh Au 1.2/1.3) that were glow discharged for 40 s using Gatan SOLARUS, blotted for 3 s under 100% humidity at 4 °C before plugged into liquid ethane cooled by liquid nitrogen using Mark IV Vitrobot (Thermo Fischer Scientific).

For all the TAS2R14-G protein complexes, cryo-EM images was collection on a Titan Krios electron microscope (Thermo Fisher Scientific) operating at 300 kV accelerating voltage, using a K3 Summit direct electron camera (Gatan) with a Gatan Quantum energy filter (operated with a slit width of 20eV). Micrographs were obtained in super-resolution mode at a pixel size of 0.832 Å and a defocus value range of -1.2 to -2.0 μm. Each video comprised 40 frames with a total dose of 60e^-^/Å^2^, exposure time was 2.0 s with a dose rate of 20e^-^/pixel/s. Automated single-particle data acquisition was collected by SerialEM software^34^ via beam tilt and astigmatism free beam-image shift^35^.

### Imaging processing

The overall cryo-EM data processing pipeline for AA_150µM_-TAS2R14-miniG_s/gust,_ AA_50µM_-TAS2R14-G_i_, AA_150µM_-TAS2R14-G_i_, FFA_1mM_-TAS2R14-miniG_s/gust_, FFA_350µM_-TAS2R14-G_i_, AA_150µM_-TAS2R14-G_g_ and apo-TAS2R14-Gi are shown in Extended Data Figs 1-3 and Extended Data Table 1. Cryo-EM movie stacks were corrected for beam-induced shifts utilizing the dose-weighting approach in Patch Motion Correction^36^. The contrast transfer function (CTF) parameters were calculated by employing the patch CTF estimation in CryoSPARC^37^.

For AA_150µM_-TAS2R14-miniG_s/gust_ complex, 13,235 movie stacks were imported in CryoSPARC v4.3. A conventional neural network-based method Topaz^38^ implemented in CryoSPARC, was used for particle picking. Total 7,013,767 particles were extracted and then subjected to 2D classification. This particle set was subjected to ab-initio reconstruction and heterogeneous refinement served as 3D classification. Then 444,009 particle projections of the best class were further applied for homogenous refinement and non-uniform refinement, generating the best density map with global resolution of 2.99 Å.

For AA_50µM_-TAS2R14-G_i_ complex, with the same steps mentioned above, a total of 10,760 movie stacks were imported in CryoSPARC v4.3. 450,989 particle projections of the best class were further applied for final homogenous refinement and Local refinement, generating the best density map with global resolution of 3.05 Å.

For AA_150µM_-TAS2R14-G_i_ complex, with the same steps mentioned above, a total of 11,241 movie stacks were imported in CryoSPARC v4.3. 851,653 particle projections of the best class were further applied for final homogenous refinement and Local refinement, generating the best density map with global resolution of 2.77 Å.

For FFA_1mM_-TAS2R14-miniG_s/gust_ complex, with the same steps mentioned above, a total of 22,659 movie stacks were imported in CryoSPARC v4.3. 224,762 particle projections of the best class were further applied for final homogenous refinement and non-uniform refinement, generating the best density map with global resolution of 3.2 Å.

For FFA_350µM_-TAS2R14-G_i_ complex, with the same steps mentioned above, a total of 9,570 movie stacks were imported in CryoSPARC v4.3. 400,545 particle projections of the best class were further applied for final homogenous refinement and Local refinement, generating the best density map with global resolution of 3.30 Å.

For AA_150µM_-TAS2R14-G_g_ complex, with the same steps mentioned above, a total of 12,041 movie stacks were imported in CryoSPARC v4.3. 243,957 particle projections of the best class were further applied for final homogenous refinement and Local refinement, generating the best density map with global resolution of 2.97 Å.

For apo-TAS2R14-G_i_ complex, with the same steps mentioned above, a total of 14,033 movie stacks were imported in CryoSPARC v4.3. 419,690 particle projections of the best class were further applied for final homogenous refinement and Local refinement, generating the best density map with global resolution of 2.94 Å.

### Model building and refinement

The coordinates of TAS2R14 from AlphaFold2 was used as the initial model. The miniG_s/gust_ heterotrimer (miniGα_s/gust_, Gβ_1_ and Gγ_2_) and Nb35 were generated using the strychnine-TAS2R46-miniG_s/gust_ complex (PDB: 7XP6) as the initial model. The G_i1_ heterotrimer (Gα_i1_, Gβ_1_ and Gγ_2_) and scFv16 were generated using the AM12033-CB2-G_i_ complex (PDB: 6KPF) as the initial model. The G_g_ heterotrimer (Gα_g_, Gβ_1_ and Gγ_2_) and scFv16 were generated using the AM12033-CB2-G_i_ complex (PDB: 6KPF) as the initial model. The cryo-EM model was fitted into the electron microscopy density map using UCSF Chimera v1.16^39^, followed by iterative manual adjustment and rebuilding in Coot v0.9.8.5^40^. Then the coordinates were refined by using Phenix^41^. The final refinement statistics are provided in Extended Data Table 1, UCSF Chimera X v1.5 were used to prepare the structural figures in the paper.

### Intracellular calcium mobilization assay

On the first day, Human Embryonic Kidney (HEK) 293T cells (ATCC, CRL-11268) were seeded at a density of one million cells per well in a six-well plate. They were then incubated overnight in a controlled cell culture environment at 37°C with 5% CO_2_ until achieving a cell confluence of 70%-75%. Subsequently, Transfection with a 1:1 DNA ratio of TAS2R14: Gα_16gust44_ (receptor or mutant 500 ng, Gα_16gust44_ 500 ng) was performed through TransIT®2020 (Mirus Biosciences). After 24 hours, cells were harvested using Versene (0.1M PBS, 0.5 mM EDTA, pH 7.4) and then seeded in a black-bottomed 384-well plate coated with poly-L-lysine at a density of 20,000 cells per well.

This was followed by an overnight incubation, after which experimental assays were conducted. The growth medium was removed, and 1× Fluo-6 calcium dye, containing 2.5mM probenecid, was added at a 20 μL per well volume. The dye was prepared in Assay buffer consisting of 1x Hank’s Balanced Salt Solution and 20 mM HEPES at pH 7.4. The plate underwent a 1-hour dark incubation at 37°C, followed by a 10-minute equilibration at room temperature before starting the readings. The baseline was established by averaging the initial ten FLIPR readings (recorded every second) prior to adding 10 μL of 3x compound. Fluorescent intensity was measured for 120 seconds following the addition of the compound. The maximum activation fluorescent intensity ratio was then calculated for each compound concentration in comparison to the baseline. The data was subsequently analyzed using GraphPad Prism 9.0 with a nonlinear regression method.

### G protein dissociation assays

G-protein bioluminescence resonance energy transfer (BRET) probes including Gα_i1_-RLuc, Gβ and Gγ-GFP and the Gα_i1_-Gβγ dissociation assay were generated and performed as previously described^42–44^. In Gα_g_-Gβγ dissociation assays, Gαg-RLuc, Gβ and Gγ-GFP were used as Gg-protein trimer dissociation BRET probes. HEK293 cells were transiently co-transfected with wild type or mutants of TAS2R14 and G-protein BRET probes. HEK293 cells were distributed into 96-well microplates at a density of 5 × 10^4^ cells per well and incubated for another 24 h at 37 °C after 24 h transfection. The cells were washed twice with Tyrode’s buffer, including 140 mM NaCl, 2.7 mM KCl, 1 mM CaCl_2_, 12 mM NaHCO_3_, 5.6 mM d-glucose, 0.5 mM MgCl_2_, 0.37 mM NaH_2_PO_4_, and 25 mM HEPES pH 7.4, and then stimulated with FFA at different concentrations as specified in the Article or figure legends. BRET signals were measured after addition of the luciferase substrate coelenterazine 400a (5 μM) using a Mithras LB940 microplate reader equipped with BRET filter sets. The BRET signal was calculated as the ratio of light emission at 510 nm/400 nm.

## Acknowledgements

This work was supported by the National Key Research and Development Program of China grant 2022YFA1302902 (T.H.), 2019YFA0904200 (J.-P.S.), the National Natural Science Foundation of China grants 32230026 (Z.-J.L.), 32271262 (T.H.), 81773704 (J.-P.S.). The Key Research Project of the Beijing Natural Science Foundation Z200019 (J.-P.S.), the CAS Strategic Priority Research Program XDB37030104 (Z.-J.L.), Shanghai Frontiers Science Center for Biomacromolecules and Precision Medicine at ShanghaiTech University. We thank the Shanghai Municipal Government and ShanghaiTech University for financial support. The cryo-EM data were collected at the Bio-Electron Microscopy Facility of ShanghaiTech University, with the assistance of L. Wang. We thank the assistance of Q.-Y. Shi at protein purification core. We thank the assistance of N. Chen and S.-W. Hu at cell expression core of iHuman Institute for their support.

## Author contribution

Z.-J. L. and T.H. initiated the project. X.-L.H. designed the expression constructs, purified the receptor complexes, prepared the final samples for cryo-EM sample and cryo-EM data collection and participated in the figure and manuscript preparation. W.-Z.A. performed the calcium mobilization functional assays assisted with Q.-W.T. M.-X.G. performed the BRET assays assisted with J.C. and F.Y. L.-J.W. EM data processing and structure determination. Y.P. participated in the EM data processing. S.-H.L. and Y.-R.W performed the molecular docking for ligands modeling. X.W. assisted with the figure preparation for G protein analysis. Q.-Q.S assisted in EM data collection. J.-L.L. and L.-Q. J. performed the insect cell expression. C.Y. participated in project discussion and edited the manuscript. J.-P.S supervised the functional assays and participating in the manuscript editing. Z.-J.L. and T.H. conceived and supervised the research, analyzed the structures and wrote the manuscript with the input from all authors.

## Data availability

The atomic coordinates for AA_150µM_-TAS2R14-miniG_s/gust,_ AA_50µM_-TAS2R14-G_i_ and AA_150µM_-TAS2R14-G_i_ have been deposited in the Protein Data Bank with the accession codes 8XQL, 8XQN and 8XQO, respectively. The EM maps for AA_150µM_-TAS2R14-miniG_s/gust,_ AA_50µM_-TAS2R14-G_i_ and AA_150µM_-TAS2R14-G_i_ have been deposited in EMDB with the codes EMD-38580, EMD-38582 and EMD-38583, respectively.

The atomic coordinates for FFA_1mM_-TAS2R14-miniG_s/gust_, and FFA_150µM_-TAS2R14-G_i_, have been deposited in the Protein Data Bank with the accession codes 8XQR and 8XQS, respectively. The EM maps for FFA_1mM_-TAS2R14-miniG_s/gust_, and FFA_150µM_-TAS2R14-G_i_, have been deposited in EMDB with the codes EMD-38586, and EMD-38587, respectively.

The atomic coordinate for AA_150µM_-TAS2R14-G_g_ and Apo-TAS2R14-G_i_ have been deposited in the Protein Data Bank with the accession codes 8XQP and 8XQT. The EM map for AA_150µM_-TAS2R14-G_g_ and Apo-TAS2R14-G_i_, have been deposited in EMDB with the code EMD-38584 and EMD-8XQT, respectively.

All other data are available upon request to the corresponding authors T.H., Z-J.L. and J.-P.S.

## Extended Data Figures

**Extended Data Fig. 1.**
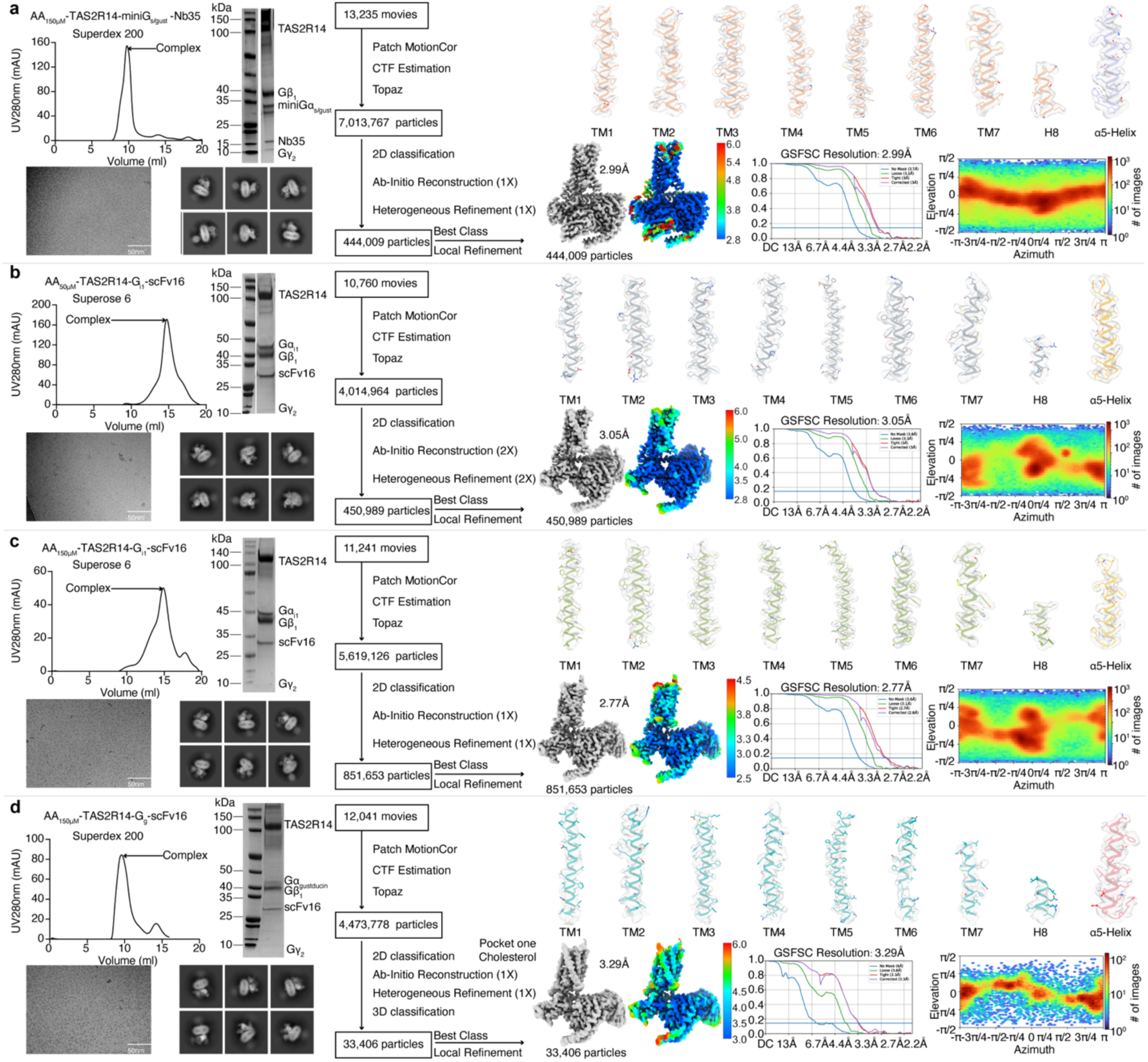
AA-TAS2R14-miniG_s/gust_-Nb35, AA-TAS2R14-G_i1_-scFv16 and AA-TAS2R14-G_g_-scFv16 sample preparation and cryo-EM data processing. **a-d**, Superdex200 size-exclusion chromatography elution profiles of the purified complex samples and SDS-PAGE analysis; Representative cryo-EM micrography (scale bar: 50 nm) and selected 2D classification showing distinct structural features of each component; Schematic representation of cryo-EM data processing workflow; Cryo-EM maps are colored by local resolution (Å); ‘Gold-standard’ FSC curve at the FSC=0.143; angular distribution of the particles used for final reconstruction; Cryo-EM maps and models of TMs for AA_150µM_-TAS2R14-miniG_s/gust_ (**a**), AA_50µM_-TAS2R14-G_i_ (**b**) and AA_150µM_-TAS2R14-G_i_ (**c**) and AA_150µM_-TAS2R14-G_g_ (**d**) complexes, respectively.

**Extended Data Fig. 2.**
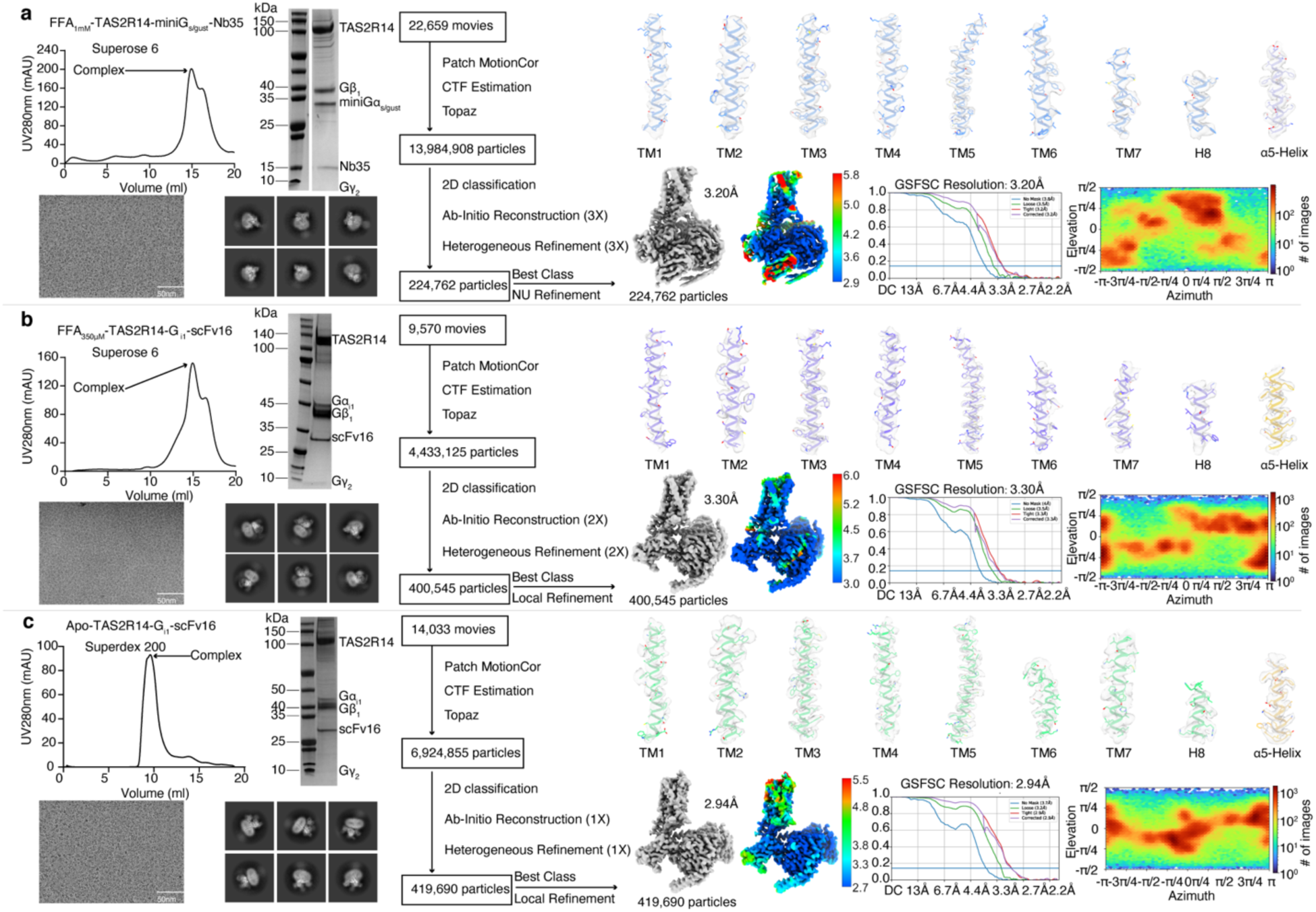
FFA-TAS2R14-miniG_s/gust_-Nb35, FFA-TAS2R14-G_i1_-scFv16 and Apo-TAS2R14-G_i1_-scFv16 sample preparation and cryo-EM data processing. **a-c,** Superdex200 size-exclusion chromatography elution profiles of the purified complex samples and SDS-PAGE analysis; Representative cryo-EM micrography (scale bar: 50 nm) and selected 2D classification showing distinct structural features of each component; Schematic representation of cryo-EM data processing workflow; Cryo-EM maps are colored by local resolution (Å); ‘Gold-standard’ FSC curve at the FSC=0.143; angular distribution of the particles used for final reconstruction; Cryo-EM maps and models of TMs for FFA_1mM_-TAS2R14-miniG_s/gust_ (**a**), FFA_350µM_-TAS2R14-G_i_ (**b**) and Apo-TAS2R14-G_i_ (**c**) complexes, respectively.

**Extended Data Fig. 3.**
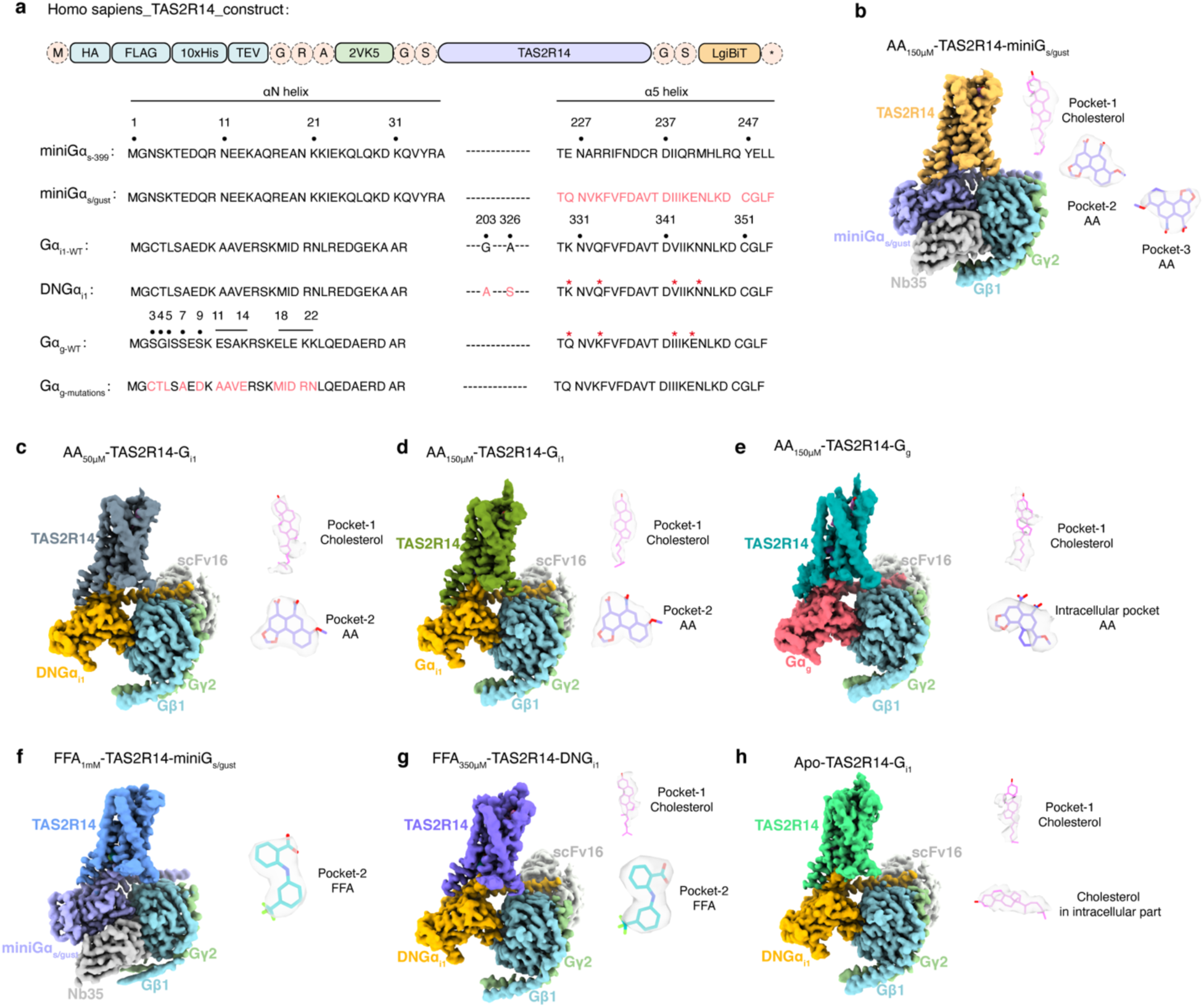
EM densities for overall structures and ligands in different complexes. **a,** Schematic representation of constructs design for the G proteins used in the complex structure determination. Red * label indicates the different residues in α5 helix between Gα_g_ and Gα_i1_. **b-h,** Cryo-EM maps of AA-bound TAS2R14 (**b-e**), FFA-bound TAS2R14 (**f-g**) and apo-TAS2R14 (**h**). The cryo-EM densities for cholesterol, AA and FFA are shown as grey meshes.

**Extended Data Fig. 4.**
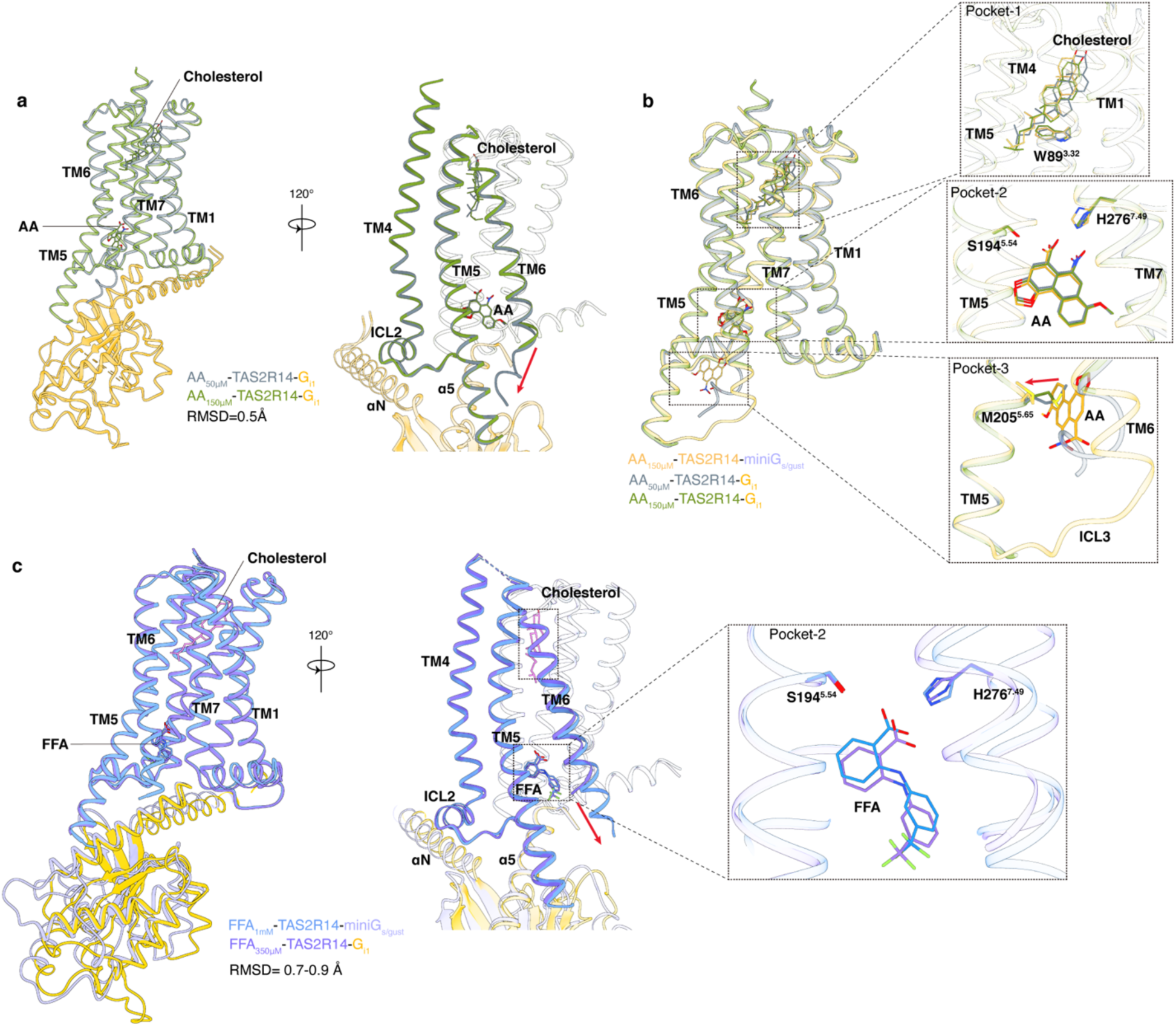
Structural comparison of AA- and FFA-bound TAS2R14 in complex with different G proteins. **a,** Structural comparison of AA_50µM_-TAS2R14-G_i1_ and AA_150µM_-TAS2R14-G_i1_ complexes. The red raw indicates the conformational differences in the cytoplasmic parts of TM6. **b,** Structural comparison of AA_150µM_-TAS2R14-miniG_s/gust_, AA_50µM_-TAS2R14-G_i1_ and AA_150µM_-TAS2R14-G_i1_ complexes. The zoom-in views of pocket-1 for cholesterol, pockets-2 and 3 for AA in the different complex structures are shown. **c,** Structural comparison of FFA_1mM_-TAS2R14-miniG_s/gust_ and FFA_350µM_-TAS2R14-G_i1_ complexes. The zoom-in view of pocket-2 for FFA in the different complex structures are shown.

**Extended Data Fig. 5.**
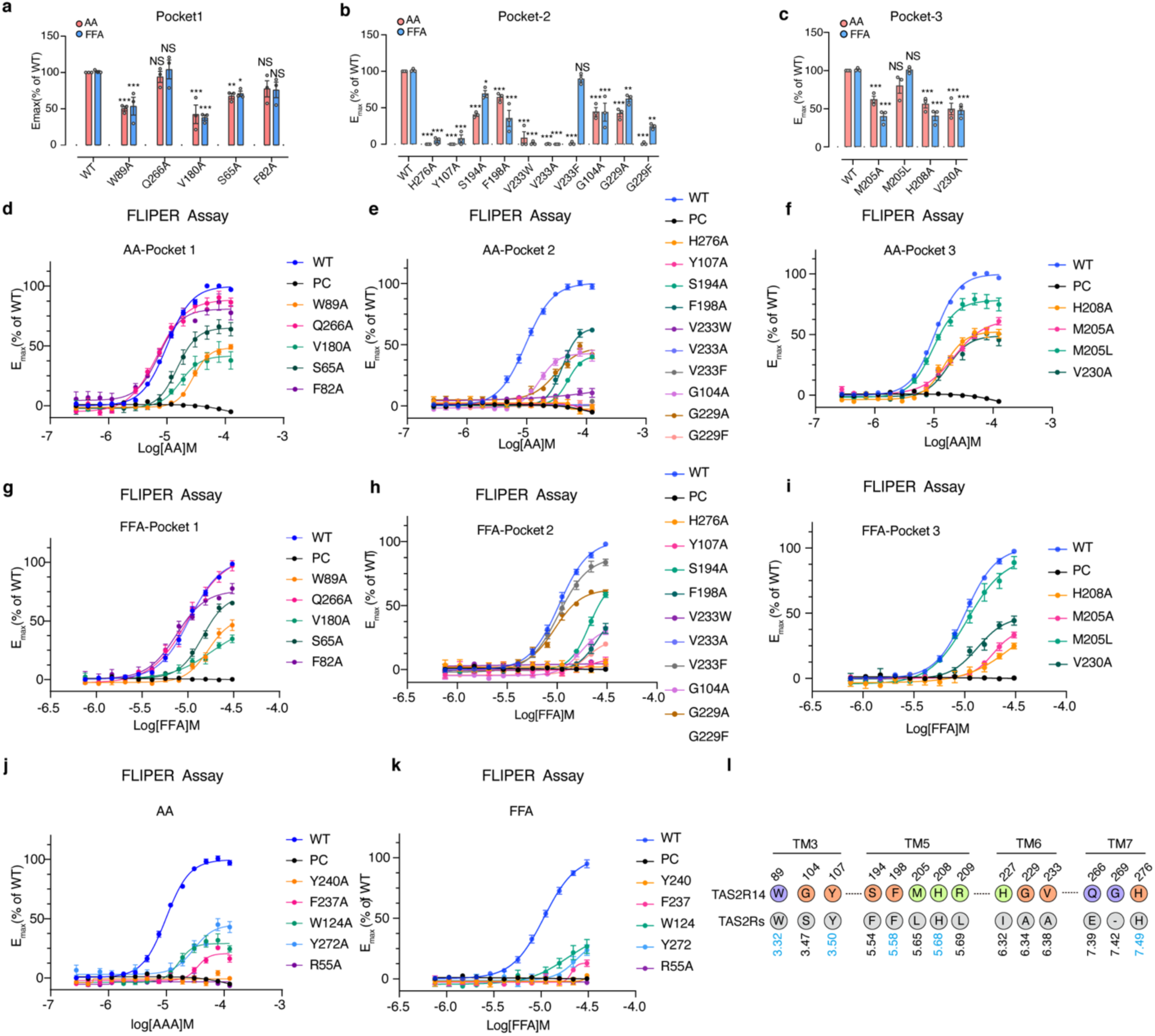
Dose responses curves of key residues mutations in ligand-binding pockets and receptor activation for AA and FFA in TAS2R14 in calcium mobilization assay. **a-c,** FLIPR Ca^2+^assay responses in wildtype (WT) TAS2R14 for mutants in pockets 1, 2 and 3 following AA or FFA stimulation. The maximum response (*E*_max_) values were calculated from the concentration–response curves (Extended Data Fig. 5). *P < 0.05; **P < 0.01; ***P < 0.001, calculated using one-way analysis of variance (ANOVA) followed by Dunnett’s test for multiple comparison analysis (with reference to the WT). NS, not significantly different between the groups. ND, signal not detectable. **d-k,** Dose-response curves of key residues mutations in calcium mobilization assay. Data are mean ± s.e.m. from 3 independent experiments (n = 3). **l,** List of key residues in pockets 1-3 in TAS2R14 and the conserved residues in TAS2Rs. The highly conserved residues are labeled in blue. Residues in purple background represent pocket-1, red for pocket-2 and green for pocket-3.

**Extended Data Fig. 6.**
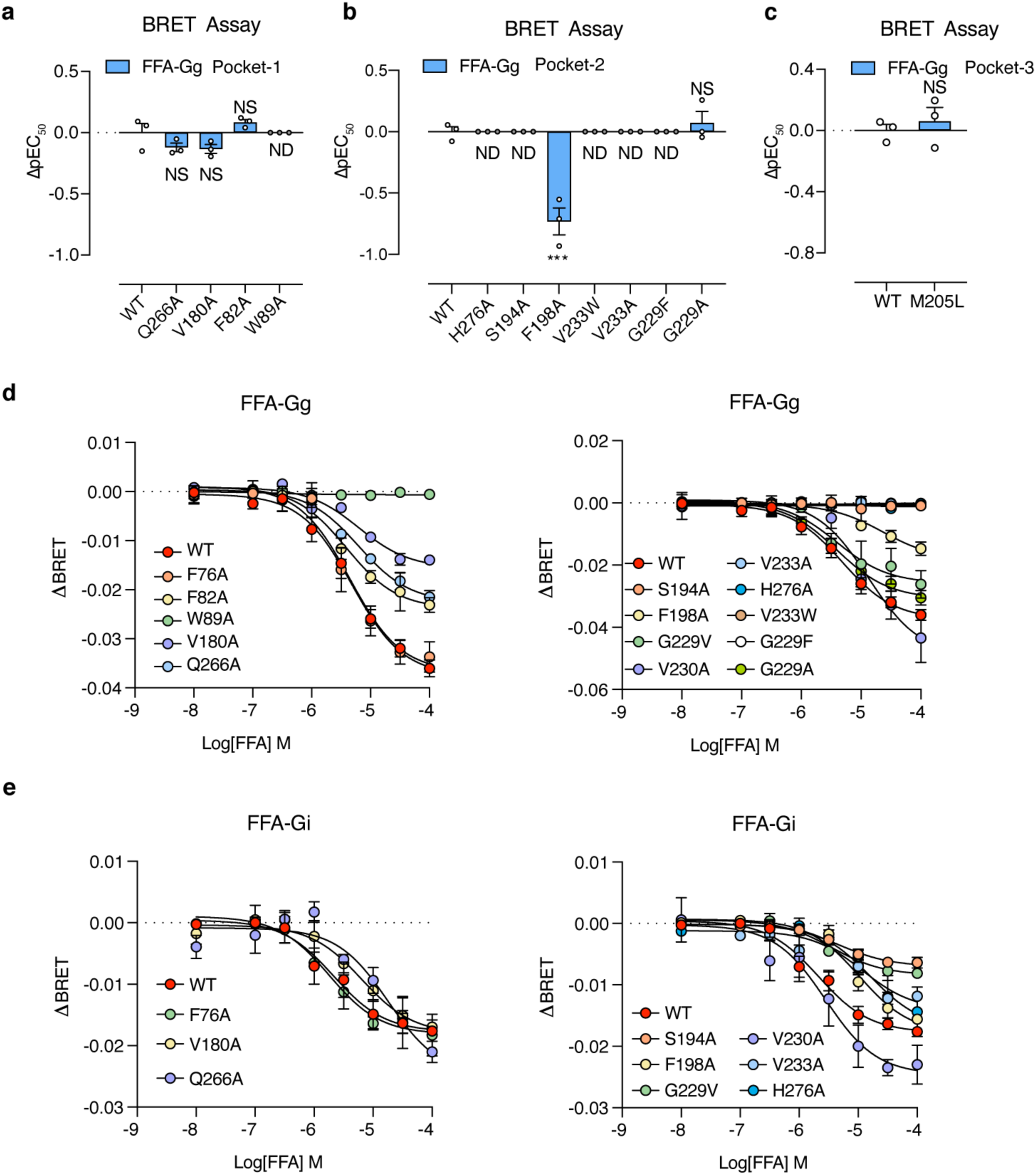
The BRET assay of FFA activated TAS2R14 in G_g_ and G_i_ pathways. **a-c,** Effects of key residue mutations in pockets 1 (**a**), 2 (**b**) and 3 (**c**) on TAS2R14’s coupling with G_g_ induced by FFA in BRET assay. Data are mean ± s.e.m. from 3 independent experiments (n = 3). Two-sided one-way ANOVA with Tukey’s test (compared with wild type). *P < 0.05; _**_P < 0.01; ***P < 0.001; NS, not significant. ND, signal not detectable. d-e, Dose response curves of key residues mutations in pockets 1, 2 and 3 on TAS2R14’s coupling with G_g_ (**d**) or G_i1_ (**e**) induced by FFA in BRET assay. Data are mean ± s.e.m. from 3 independent experiments (n = 3).

**Extended Data Fig. 7.**
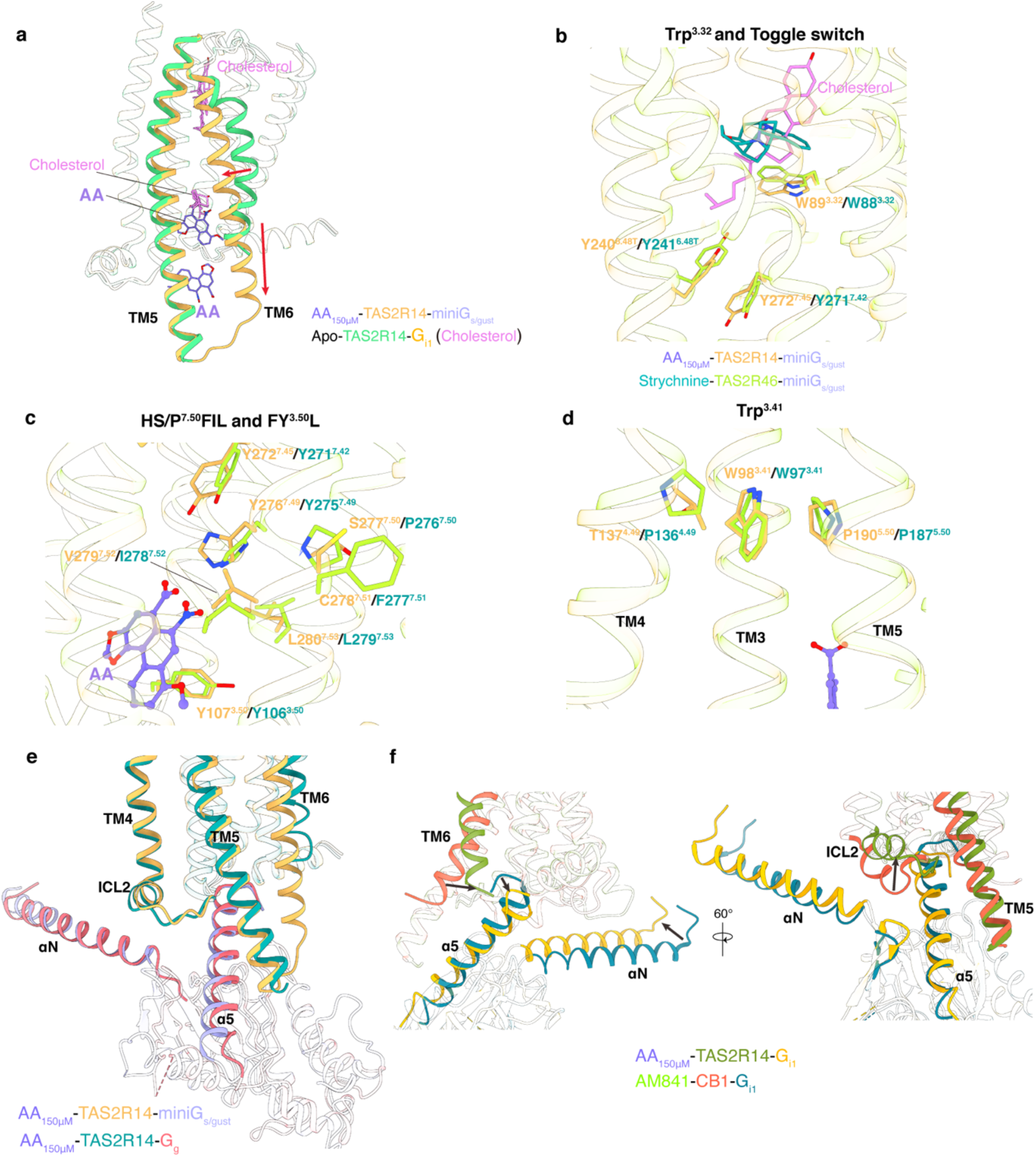
Structural comparison of active features and G protein coupling in TAS2R14. **a,** Structural comparison of receptors in AA_150µM_-TAS2R14-miniG_s/gust_ and Apo-TAS2R14-G_i_ complexes. TM5 and TM6 show different conformational changes. The red raw indicates the extension of cytoplasmic parts of TM6 after AA binding to pockets 2 and 3. **b-d,** Structural comparison of key motifs related to activation in TAS2R14 and TAS2R46. Trp^3.32^ and toggle switch residues (**b**), HS/P^7.50^FIL and FY^3.50^L (**c**) and Trp^3.41^ residue (**d**), respectively. e, Structure comparison of the miniGα_s/gust_ and Gα_g_ binding modes in TAS2R14. AA_150µM_-TAS2R14-miniG_s/gust_ and AA_150µM_-TAS2R14-G_g_ complex structures are used for analysis. **f,** Structure comparison of the Gα_i_ binding modes in TAS2R14 and CB1.

**Extended Data Table 1.**
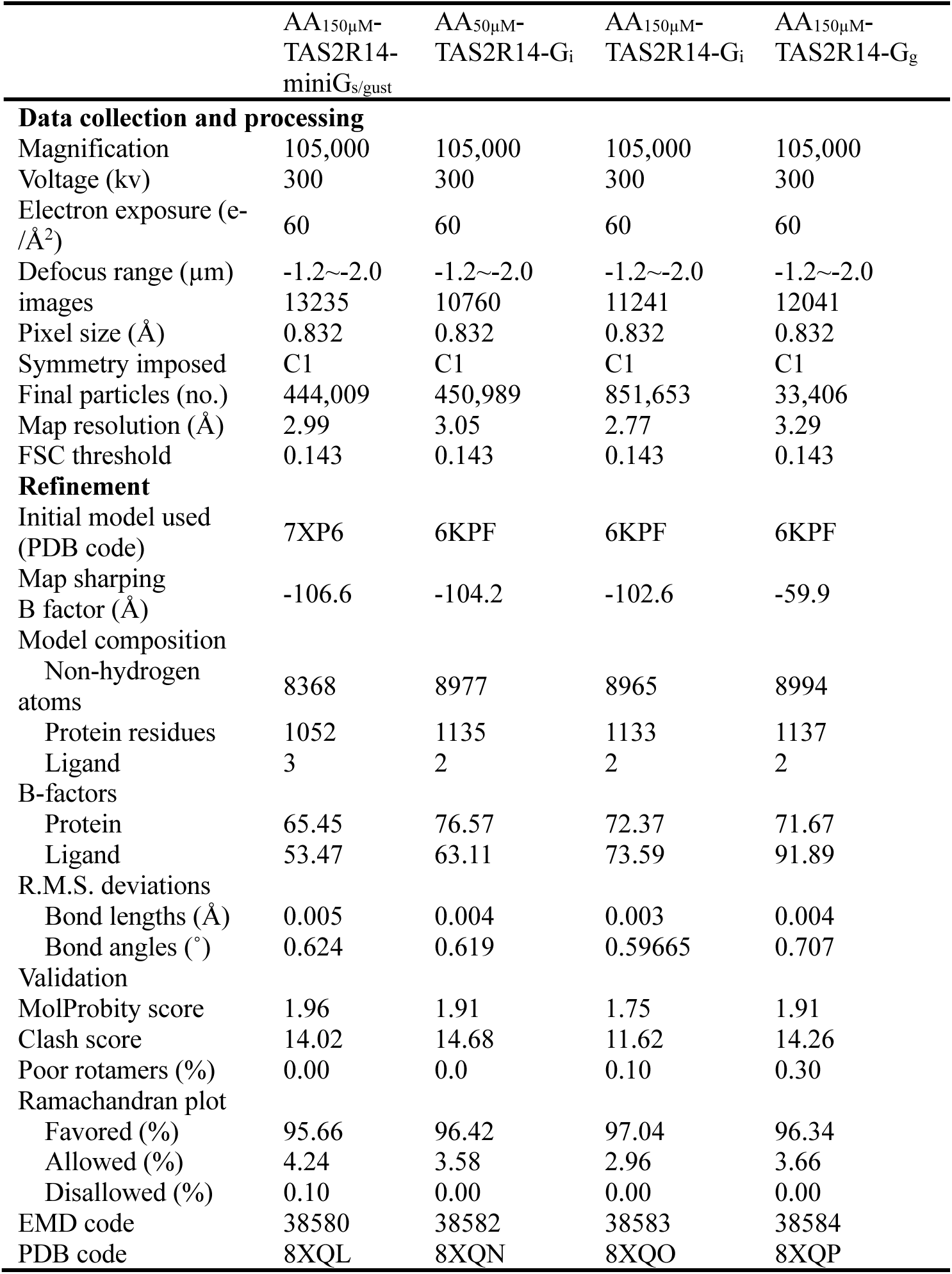
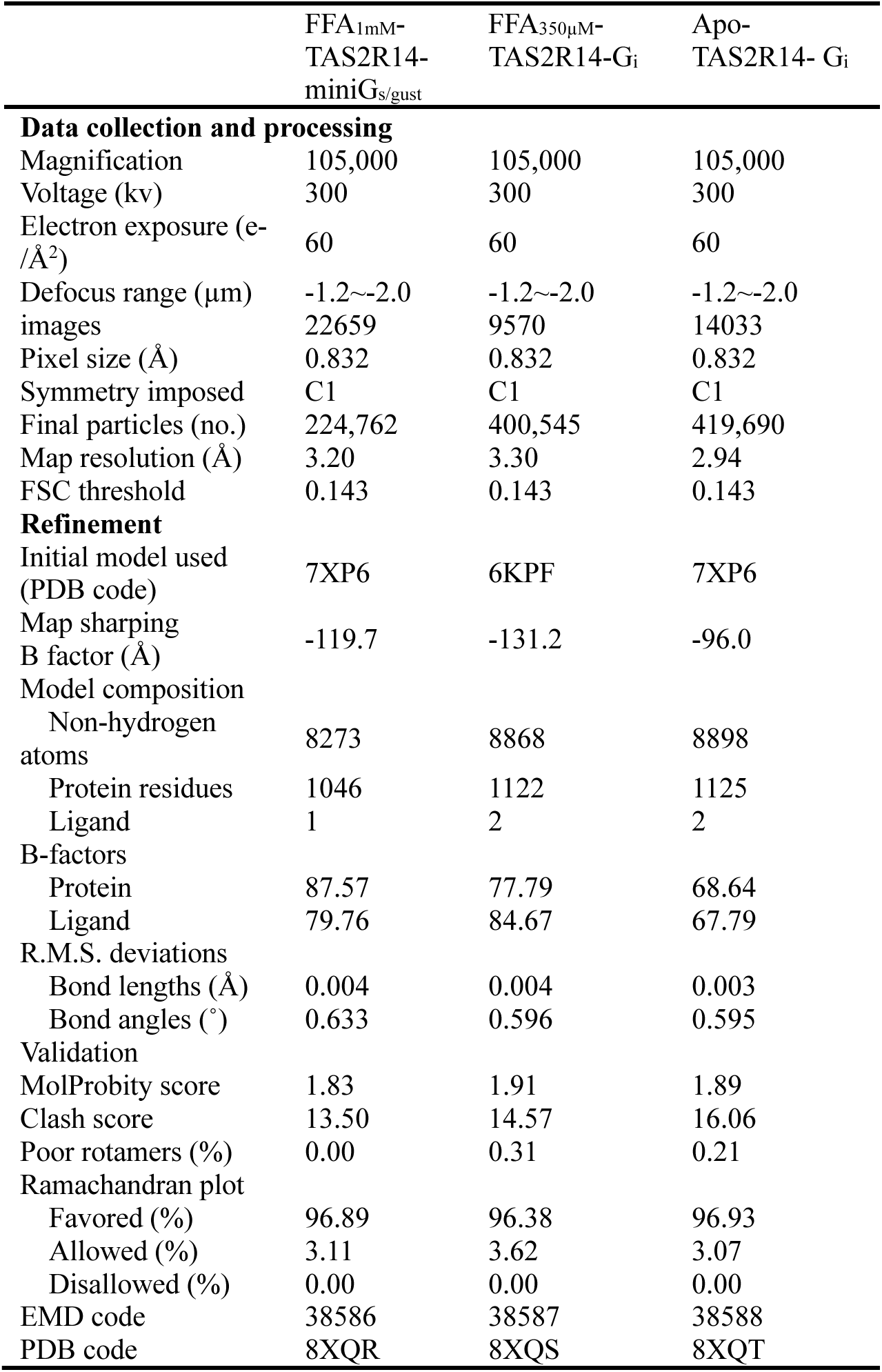
Cryo-EM data collection, model refinement, and validation statistics.

**Extended Data Table 2.** Activity of AA or FFA on wild-type TAS2R14 and mutants measured by calcium mobilization assay.

| Mutation | pEC<sub>50</sub> ± s.e.m.<sup>#</sup> | E<sub>max</sub> | P value | N | pEC<sub>50</sub> ± s.e.m.<sup>#</sup> | E<sub>max</sub> | P value | N | Expression (%WT)<sup>\$</sup> |
| --- | --- | --- | --- | --- | --- | --- | --- | --- | --- |
| AA |  |  |  |  | FFA |  |  |  |  |
| WT | 4.98±0.02 | 100 |  | 3 | 4.99±0.04 | 100 |  | 3 | 100 |
| W89A | 4.54±0.04 | 50.4±3.2 | *** | 3 | 4.83±0.02 | 59.67±7.21 | *** | 3 | 122.6±11.4 |
| Q266A | 5.17±0.04 | 93.97±7.59 | NS | 3 | 5.07±0.05 | 104.1±12.9 | NS | 3 | 113.1±10.1 |
| V180A | 4.79±0.09 | 62.7±12.14 | *** | 3 | 4.81±0.02 | 32.41±4.99 | *** | 3 | 131.1±15.8 |
| S65A | 4.82±0.02 | 67.48±3.86 | ** | 3 | 4.84±0.02 | 70.74±3.6 | * | 3 | 155.6±8 |
| F82A | 5.1±0.04 | 80.93±8.25 | NS | 3 | 5.1±0.01 | 75.21±11.3 | NS | 3 | 127.1±24.2 |
| H276A | ND | ND | ND | 3 | ND | ND | ND | 3 | 115.9±23.1 |
| Y107A | ND | ND | ND | 3 | ND | ND | ND | 3 | 102.4±11.5 |
| S194A | 4.31±0.01 | 40.39±2.17 | *** | 3 | 4.68±0.03 | 69.13±4.44 | * | 3 | 100.9±5.6 |
| F198A | 4.39±0.06 | 64.39±3.59 | *** | 3 | 4.58±0.08 | 35.46±11.0 | *** | 3 | 123.7±7.5 |
| V233W | ND | ND | ND | 3 | ND | ND | ND | 3 | 85.81±18.9 |
| V233A | ND | ND | ND | 3 | ND | ND | ND | 3 | 97.74±13.2 |
| V233F | ND | ND | ND | 3 | 4.99±0.03 | 89.54±4.15 | NS | 3 | 106.3±19.7 |
| G104A | 4.79±0.04 | 44.48±5.89 | *** | 3 | 4.73±0.06 | 43.94±12.3 | *** | 3 | 143±7.8 |
| G229A | 4.53±0.08 | 42.12±4.3 | *** | 3 | 5.02±0.02 | 62.09±4.03 | ** | 3 | 103.2±17.8 |
| G229F | ND | ND | ND | 3 | 4.76±0.04 | 24.4±2.21 | *** | 3 | 129.9±7.1 |
| M205A | 4.67±0.06 | 62±4.43 | *** | 3 | 4.71±0.06 | 39.91±5.34 | *** | 3 | 81.67±15 |
| M205L | 5.02±0.03 | 80.01±9.68 | NS | 3 | 4.96±0.03 | 100.7±2.45 | NS | 3 | 106.9±12.6 |
| H208A | 4.83±0.03 | 56.23±5.2 | *** | 3 | 4.67±0.01 | 40.28±5.8 | *** | 3 | 84.94±10.4 |
| V230A | 4.78±0.04 | 49.69±7.6 | *** | 3 | 4.89±0.07 | 47.52±5.25 | *** | 3 | 135.9±73.5 |
| Y240A | ND | ND | ND | 3 | ND | ND | ND | 3 | 112.8±9.5 |
| F237A | 4.43±0.07 | 22.89±5.23 | *** | 3 | 4.7±0.04 | 22.32±6.63 | *** | 3 | 121.3±19.4 |
| W124A | 4.79±0.01 | 32.82±7.89 | *** | 3 | 4.64±0.12 | 50.03±9.61 | *** | 3 | 146.1±13.6 |
| Y272A | 4.58±0.03 | 33.55±8.22 | *** | 3 | 4.69±0.06 | 32.75±7.12 | *** | 3 | 105.4±20.6 |
| R55A | ND | ND | ND | 3 | ND | ND | ND | 3 | 104.3±23.1 |

<sup>#</sup>Data are mean ± s.e.m. from at least three independent experiments. \**p*<0.05, \*\**p*<0.01, \*\*\**p*<0.001 by one-way analysis of variance followed by Dunnett's post-test compared to the response of wild type. NS, not significant; ND, signal not detectable.
<sup>\$</sup>Protein expression levels of TAS2R14 constructs at the cell surface were determined in parallel by flow cytometry with an anti-Flag antibody (Sigma) and reported as per cent compared to the wild type from three independent measurements performed in technical duplicate.

## References

1 Waterloo, L. et al. Discovery of 2-Aminopyrimidines as Potent Agonists for the Bitter Taste Receptor TAS2R14. J Med Chem 66, 3499–3521 (2023). 10.1021/acs.jmedchem.2c01997

2 Levit, A. et al. The bitter pill: clinical drugs that activate the human bitter taste receptor TAS2R14. FASEB J 28, 1181–1197 (2014). 10.1096/fj.13-242594

3 Fierro, F. et al. Inhibiting a promiscuous GPCR: iterative discovery of bitter taste receptor ligands. Cell Mol Life Sci 80, 114 (2023). 10.1007/s00018-023-04765-0

4 Deshpande, D. A. et al. Bitter taste receptors on airway smooth muscle bronchodilate by localized calcium signaling and reverse obstruction. Nat Med 16, 1299–1304 (2010). 10.1038/nm.2237

5 Shaik, F. A., Medapati, M. R. & Chelikani, P. Cholesterol modulates the signaling of chemosensory bitter taste receptor T2R14 in human airway cells. Am J Physiol Lung Cell Mol Physiol 316, L45–L57 (2019). 10.1152/ajplung.00169.2018

6 Kooistra, A. J. et al. GPCRdb in 2021: integrating GPCR sequence, structure and function. Nucleic Acids Res 49, D335–D343 (2021). 10.1093/nar/gkaa1080

7 Lundstrom, J. N., Boesveldt, S. & Albrecht, J. Central Processing of the Chemical Senses: an Overview. ACS Chem Neurosci 2, 5–16 (2011). 10.1021/cn1000843

8 Chandrashekar, J., Hoon, M. A., Ryba, N. J. & Zuker, C. S. The receptors and cells for mammalian taste. Nature 444, 288–294 (2006). 10.1038/nature05401

9 Lee, S. J., Depoortere, I. & Hatt, H. Therapeutic potential of ectopic olfactory and taste receptors. Nat Rev Drug Discov 18, 116–138 (2019). 10.1038/s41573-018-0002-3

10 Clark, A. A., Liggett, S. B. & Munger, S. D. Extraoral bitter taste receptors as mediators of off-target drug effects. FASEB J 26, 4827–4831 (2012). 10.1096/fj.12-215087

11 Dagan-Wiener, A. et al. BitterDB: taste ligands and receptors database in 2019. Nucleic Acids Res 47, D1179–D1185 (2019). 10.1093/nar/gky974

12 Bloxham, C. J., Foster, S. R. & Thomas, W. G. A Bitter Taste in Your Heart. Front Physiol 11, 431 (2020). 10.3389/fphys.2020.00431

13 Kim, D., Pauer, S. H., Yong, H. M., An, S. S. & Liggett, S. B. beta2-Adrenergic Receptors Chaperone Trapped Bitter Taste Receptor 14 to the Cell Surface as a Heterodimer and Exert Unidirectional Desensitization of Taste Receptor Function. J Biol Chem 291, 17616–17628 (2016). 10.1074/jbc.M116.722736

14 Nayak, A. P., Shah, S. D., Michael, J. V. & Deshpande, D. A. Bitter Taste Receptors for Asthma Therapeutics. Front Physiol 10, 884 (2019). 10.3389/fphys.2019.00884

15 Woo, J. A. et al. A Par3/LIM Kinase/Cofilin Pathway Mediates Human Airway Smooth Muscle Relaxation by TAS2R14. Am J Respir Cell Mol Biol 68, 417–429 (2023). 10.1165/rcmb.2022-0303OC

16 Wong, G. T., Gannon, K. S. & Margolskee, R. F. Transduction of bitter and sweet taste by gustducin. Nature 381, 796–800 (1996). 10.1038/381796a0

17 Ming, D., Ruiz-Avila, L. & Margolskee, R. F. Characterization and solubilization of bitter-responsive receptors that couple to gustducin. Proc Natl Acad Sci U S A 95, 8933–8938 (1998). 10.1073/pnas.95.15.8933

18 Kim, D., Woo, J. A., Geffken, E., An, S. S. & Liggett, S. B. Coupling of Airway Smooth Muscle Bitter Taste Receptors to Intracellular Signaling and Relaxation Is via G(alphai1,2,3). Am J Respir Cell Mol Biol 56, 762–771 (2017). 10.1165/rcmb.2016-0373OC

19 Sainz, E. et al. Functional characterization of human bitter taste receptors. Biochem J 403, 537–543 (2007). 10.1042/BJ20061744

20 Winder, C. V., Wax, J., Serrano, B., Jones, E. M. & Mc, P. M. Anti-inflammatory and antipyretic properties of N-(alpha,alpha,alpha-trifluoro-m-tolyl) anthranilic acid (CI-440; flufenamic acid). Arthritis Rheum 6, 36–47 (1963). 10.1002/art.1780060105

21 Wooding, S. P. et al. Association of a bitter taste receptor mutation with Balkan Endemic Nephropathy (BEN). BMC Med Genet 13, 96 (2012). 10.1186/1471-2350-13-96

22 Wang, Z. X. et al. Discovery of TAS2R14 Agonists from Platycodon grandiflorum Using Virtual Screening and Affinity Screening Based on a Novel TAS2R14-Functionalized HEMT Sensor Combined with UPLC-MS Analysis. J Agric Food Chem 66, 11663–11671 (2018). 10.1021/acs.jafc.8b04455

23 Martin, L. T. P. et al. Bitter taste receptors are expressed in human epithelial ovarian and prostate cancers cells and noscapine stimulation impacts cell survival. Mol Cell Biochem 454, 203–214 (2019). 10.1007/s11010-018-3464-z

24 Xu, W. et al. Structural basis for strychnine activation of human bitter taste receptor TAS2R46. Science 377, 1298–1304 (2022). 10.1126/science.abo1633

25 Nowak, S. et al. Reengineering the ligand sensitivity of the broadly tuned human bitter taste receptor TAS2R14. Biochim Biophys Acta Gen Subj 1862, 2162–2173 (2018). 10.1016/j.bbagen.2018.07.009

26 Topin, J. et al. Functional molecular switches of mammalian G protein-coupled bitter-taste receptors. Cell Mol Life Sci 78, 7605–7615 (2021). 10.1007/s00018-021-03968-7

27 Dixon, A. S. et al. NanoLuc Complementation Reporter Optimized for Accurate Measurement of Protein Interactions in Cells. Acs Chem Biol 11, 400–408 (2016). 10.1021/acschembio.5b00753

28 Duan, J. et al. NanoBit. Nat Commun 11 (2020). https://doi.org/ARTN 4121 10.1038/s41467-020-17933-8

29 Liu, P. et al. The structural basis of the dominant negative phenotype of the Gα βγG203A/A326S heterotrimer. Acta Pharmacol Sin 37, 1259–1272 (2016). 10.1038/aps.2016.69

30 Ammon, C., Schäfer, J., Kreuzer, O. J. & Meyerhof, W. Presence of a plasma membrane targeting sequence in the amino-terminal region of the rat somatostatin receptor 3. Arch Physiol Biochem 110, 137–145 (2002). 10.1076/apab.110.1.137.908

31 Ueda, T., Ugawa, S., Yamamura, H., Imaizumi, Y. & Shimada, S. G16g44. J Neurosci 23, 7376–7380 (2003).

32 Rasmussen, S. G. et al. Crystal structure of the beta2 adrenergic receptor-Gs protein complex. Nature 477, 549–555 (2011). 10.1038/nature10361

33 Hua, T. et al. Activation and Signaling Mechanism Revealed by Cannabinoid Receptor-G(i) Complex Structures. Cell 180, 655–665 e618 (2020). 10.1016/j.cell.2020.01.008

34 Mastronarde, D. N. SerialEM. J Struct Biol 152, 36–51 (2005). 10.1016/j.jsb.2005.07.007

35 Wu, C., Huang, X., Cheng, J., Zhu, D. & Zhang, X. High-quality, high-throughput cryo-electron microscopy data collection via beam tilt and astigmatism-free beam-image shift. J Struct Biol 208, 107396 (2019). 10.1016/j.jsb.2019.09.013

36 Zheng, S. Q. et al. MotionCor2: anisotropic correction of beam-induced motion for improved cryo-electron microscopy. Nat Methods 14, 331–332 (2017). 10.1038/nmeth.4193

37 Punjani, A., Rubinstein, J. L., Fleet, D. J. & Brubaker, M. A. cryoSPARC: algorithms for rapid unsupervised cryo-EM structure determination. Nat Methods 14, 290-+ (2017). 10.1038/Nmeth.4169

38 Bepler, T. et al. Positive-unlabeled convolutional neural networks for particle picking in cryo-electron micrographs. Nat Methods 16, 1153–1160 (2019). 10.1038/s41592-019-0575-8

39 Pettersen, E. F. et al. UCSF Chimera--a visualization system for exploratory research and analysis. J Comput Chem 25, 1605–1612 (2004). 10.1002/jcc.20084

40 Emsley, P. & Cowtan, K. Coot: model-building tools for molecular graphics. Acta Crystallogr D Biol Crystallogr 60, 2126–2132 (2004). 10.1107/S0907444904019158

41 Adams, P. D. et al. Recent developments in the PHENIX software for automated crystallographic structure determination. J Synchrotron Radiat 11, 53–55 (2004). 10.1107/s0909049503024130

42 Guo, L. et al. Structural basis of amine odorant perception by a mammal olfactory receptor. Nature 618, 193–200 (2023). 10.1038/s41586-023-06106-4

43 Cheng, J. et al. Autonomous sensing of the insulin peptide by an olfactory G protein-coupled receptor modulates glucose metabolism. Cell Metab 34, 240–255 e210 (2022). 10.1016/j.cmet.2021.12.022

44 Olsen, R. H. J. et al. TRUPATH, an open-source biosensor platform for interrogating the GPCR transducerome. Nat Chem Biol 16, 841–849 (2020). 10.1038/s41589-020-0535-8

